# A Combinatorial Approach of Biparental QTL Mapping and Genome-Wide Association Analysis Identifies Candidate Genes for Phytophthora Blight Resistance in Sesame

**DOI:** 10.1101/2020.03.18.996637

**Authors:** Asekova Sovetgul, Eunyoung Oh, Krishnanand P. Kulkarni, Myoung Hee Lee, Jung In Kim, Suk-Bok Pae, Minsu Kim, Ki-Won Oh, Kwang-Soo Cho, Sungup Kim

## Abstract

Phytophthora blight, caused by pathogen *Phytophthora nicotianae*, is responsible for a huge reduction in sesame (*Sesamum indicum* L.) crop yields. In this study, we utilized a combinatorial approach involving biparental QTL mapping and genome-wide association (GWAS) analysis to identify genes associated with Phytophthora blight resistance in sesame. Evaluation of resistant of the parental varieties (Goenbaek, Osan and Milsung) and the RILs of both the populations in greenhouse conditions suggested the qualitative nature of the trait.. The genetic map comprised thirteen LGs covering a total map length of 887.49 cM with an average inter-marker distance of 4.69 cM. Significant QTLs explaining phenotypic variation in the range of 2.25% to 69.24% were identified on chromosomes 10 and 13 (Chr10 and Chr13). A resistance locus detected on Chr10 was found to be highly significant. The association of this locus to PBR was also identified through BSA and single marker analysis in Goenbaek × Milsung cross and through genome-wide association mapping of 87 sesame accessions. The GWAS analysis identified 44 SNP loci significantly associated with Phytophthora disease-resistant traits on Chr10. Further, the haplotype block analysis conducted in order to find whether the SNPs associated with resistance in this study showed that the SNPs are in high LD with the resistance QTL. We obtained a total of 68 candidate genes, which included a number of defense-related *R* genes. One of the genes, *SIN_1019016* (*At1g58390)* showed high expression in the resistant parent. The results from this study would be highly useful in identifying genetic and molecular factors associated with Phytophthora blight resistance in sesame.

## 1 Introduction

Sesame (*Sesamum indicum* L.) (2n = 2x = 26), comprising a genome of 357 Mbp arranged to 13 linkage groups, is one of the orphan crops with limited genomic resources (Wang et al., 2016). Among the edible oilseed crops, sesame occupies a unique position as it can be cultivated throughout the year and its polyunsaturated fatty acid (PUFA) content makes it beneficial for human health. It is rich in oil (52-53%) and protein (26.25%). The oil extracted from the tiny seeds is odorless, very stable, and contains an antioxidant system comprising of sesamol and sesamolinol formed from sesamolin, which substantially reduces its oxidation rate. Sesame oil is reported to be effective in reducing stress and tension and in preventing nervous disorders, various types of cancers, relieving fatigue and promoting strength and vitality (Majdalawieh et al., 2017; Nakimi, 1995). The global demand for vegetable oils is increasing and estimated to touch 240 million tons by 2050 (Barcelos et al., 2015). Sesame is, therefore, a productive plant that may greatly contribute to meet this demand. However, sesame growth and production have a number of serious difficulties such as shattering, lodging, indeterminate growth, waterlogging stress together with diseases such as phytophthora blight, wilt, phyllody, powdery mildew, bacterial blight and charcoal problem causing significant yield losses every year. Among these, Phytophthora blight is one of the important biotic factors for intensive management of sesame crops in countries worldwide having excessive rainfall and humid environments.

Phytophthora blight is one of the most damaging diseases of sesame worldwide. It is caused by *Phytophthora nicotianae* Breda de Haan and occurs in sesame growing regions with moist and humid conditions. *P. nicotianae* has been found to be the dominant species in Brazil, Egypt, South Africa, and Tunisia (Panabières et al., 2016). The optimum temperature for its growth and infection affecting plants is between 33°C to 36°C and may be considered as an emerging pathogen in the current context of global climate change. In Korean Peninsula, it has been reported as the most important soil born disease causing severe economic losses (Choi et al., 1987). Unfortunately, little is known on the existence of reliable sources of resistance, and very few efforts have been made to develop high-yielding cultivars having resistance to Phytophthora blight. The control measures applied to affected fields are often impractical and expensive.

Considerable variability in reactions of sesame germplasm to *P. nicotianae* has been reported worldwide and several research groups have done germplasm evaluation studies to select for the resistant varieties (Oh et al., 2018). So far, the Korean National Agrobiodiversity Germplasm has identified four *P. nicotianae* isolates, namely ‘KACC48120’, ‘48121’, ‘No2526’ and ‘2040’ (Oh et al., 2018). Oh et al. (2018) evaluated sesame genotypes for *P. nicotianae* disease resistance in greenhouse nursery that were artificially inoculated with four strains, namely, KACC48120, 48121, No2526 and 2040. These strains were collected from infected sesame plants in Gyeongju, Naju, Gunwi, and Yecheon province fields in South Korea. To control the damage caused by Phytophthora blight, the application of plant resistance through breeding programs are considered the most effective method. Most commercial cultivars and populations of *Sesamum* spp. lack sufficient genetic resistance to avoid fungicide use for Phytophthora blight, and there is no information on the inheritance of resistance to the disease in this species in sesame. Hence, there is a need to assess the genetic diversity among the present sesame cultivars and evaluate them for the resistance to *P. nicotianae*. The genetic diversity and population structure analysis may help identify the genetically divergent varieties which can be used for breeding purpose (Asekova et al., 2018). Such sources are used to identify the genetic loci (QTLs or markers linked to the trait) responsible for the resistance to *P. nicotianae*.

A large number of molecular markers such as simple sequence repeat (SSR) and single-nucleotide polymorphism (SNP) markers have been made available in sesame due to the recent developments in next-generation sequencing (NGS) technologies. These markers can be utilized for diversity analysis as well as to develop high-density genetic linkage maps, which can be later used to identify the QTLs associated with a trait of interest in a limited number of crosses. Further, genome-wide association analysis (GWAS) can be performed by screening a large number of germplasm with these markers to identify marker-trait associations, which may find useful in marker-assisted breeding for variety development.

SNP markers have become increasingly important tools for molecular genetic analysis, as single base-pair changes are the most abundant small-scale genetic variation present between related sequences of DNA. The SNPs are stable, heritable, and abundant in the plant genomes and are therefore preferred choice of markers for linkage mapping and genome-wide association studies (Varshney et al., 2012; Zhang et al., 2016). Alternatively, SSRs are also widely used in genetic analyses because of their locus specificity, multi-allelic and codominant nature (Tautz, 1989). Using these and other markers systems, several interspecific and intraspecific genetic linkage maps have been developed in sesame (Mei et. al., 2017; Liu et al., 2015; Wei et. al., 2009; Wang et. al., 2017). The first intraspecific linkage map (Wei et al., 2009) was constituted of 8 EST-SSR (Expressed Sequence Tags), 25 AFLP (Amplified Fragment Length Polymorphism) and 187 RSAMPL (Random Selective Amplification of Microsatellite Polymorphic Loci) markers were grouped into 30 linkage groups (LGs) spanning 936.72 cM. Subsequently, two intraspecific genetic maps were constructed using backcross (BC_1_) by (Specific Length Amplified Fragment Sequencing (SLAF) and recombinant inbred line (RIL) population using 424 SSR markers grouped into 13 actual sesame linkage groups (SLG) (Mei et al., 2017; Wang et al., 2017). Nevertheless, the level of marker saturation is still low in the context of both fine mapping and genomic synteny. As an orphan or neglected crop, a high-density genetic map in sesame is still limited. Six dense SNP genetic maps have been so far constructed for sesame (Mei et al., 2017; Uncu et al., 2016; Wu et al., 2014; Wang et al., 2016; Zhang et al., 2013a; 2016). Most of these six genetic maps have been applied for QTL mapping using reduced representation genome sequencing (RRGS), while only the genotyping-by-sequencing (GBS) generated 770 SNPs and 50 SSRs were used to construct the first sesame linkage map (Uncu et al., 2016). Moreover, SSR markers widely used for genetic diversity and association mapping of germplasm lines to analyze quantitative traits or tagging to the various disease-related genes (Li et al., 2014; Liu et al., 2015; Wu et al., 2014; Wang et al., 2017). Four main complexity reduction methods have been described to date: deep resequencing according to the principal of paired-end sequencing and *de novo* sequencing (RNA-Seq) using Illumina HiSeq 2000 platform can detect SNPs, small InDels and structural variants, restriction site-associated DNA (RAD) sequencing, reduced representation libraries (RRL), specific-locus amplified fragment (SLAF) sequencing, and GBS. Using the SLAF approach in sesame, 89,924, 71,793, and 9378 in three different studies (Cui et al., 2017; Mei et al., 2017; Zhang et al., 2013b). Recently, RAD sequencing has been applied in sesame for fine mapping of plant height and seed coat color (Wang et al., 2016).

In the present study, we performed a genome-wide association study (GWAS) using GBS-derived SNPs in a set of sesame germplasm to identify the chromosomal regions associated with Phytophthora blight resistance. Further, we selected 1 resistant and two susceptible parents from the GWAS sesame panel to develop two bi-parental mapping populations, which were later genotyped using GBS to identify the QTL associated with Phytophthora blight resistance.

## Results

### Phenotype evaluation of RIL populations and germplasm set

In this study, Phytophthora blight resistance of the parental varieties (Goenbaek, Osan, and Milsung) and the RILs of both the populations were evaluated in greenhouse conditions. Based on the standard evaluation method, the plants (F_2_) of cross between Goenbaek and Osan, segregated phenotypically as 128 resistant and 333 susceptible lines, fitting the expected phenotypic ratio of 1:3 with χ^2^=1.88 and *P*=0.18 (*P*>0.05) (Table S2). For the Goenbaek and Milsung F2 plants, the segregation ratio also followed the expected segregation ratio for a recessive gene of 1:3 (R:S), with χ^2^=2.07 and *P*=0.16 (*P*>0.05). (Table S2). This findings indicative of a monogenic and recessive control involving a single copy of resistance allele in Goenbaek. From the Population-I, 47 RILs showed resistance, whereas 43 RILs were found to be susceptible to *P. nicotianae* isolate KACC48121 (Table 1). Similarly, from Population-II, 90 RILs were resistant whereas 98 RILs were found to be susceptible to *P. nicotianae* isolate KACC48121. Both the RIL populations showed 1:1 ratio of resistant:susceptible plants (*P* < *0*.*05* and *P* < *0*.*01*), suggesting the qualitative behavior of the Phytophthora blight (Table 1). The frequency distribution of traits observed was race-specific qualitative and are presented in Figures 1 and 2.

**Table 1.** Phenotypic evaluation information and SSR marker segregation in the RIL populations.

| Isolate | Phenotype |  |  |  |  |  |
| --- | --- | --- | --- | --- | --- | --- |
|  | Population-I (G□O) |  |  | Population-II (G□M) |  |  |
| | No. of resistant plants | No. of susceptible plants | $\chi^2$ | No. of resistant plants | No. of susceptible plants | $\chi^2$ |
| KACC48121 | 47 | 43 | 0.178 | 90 | 98 | 0.340 |
| Marker | Genotype |  |  |  |  |  |
|  | Population-I (G□O) |  |  | Population-II (G□M) |  |  |
| | RR=R | rr=S | $\chi^2(1:1)^a$ | RR=R | rr=S | $\chi^2(1:1)^a$ |
| SiSSM85570 | — | — | — | 95 | 93 | 0.020ns |
| SiSSM85354 | — | — | — | 89 | 95 | 0.200ns |
| SiSSM84996 | 42 | 46 | 0.180ns | — | — | — |
| SiSSM84917 | 46 | 43 | 0.100ns | 90 | 94 | 0.090ns |
| SiSSM85023 | 46 | 43 | 0.100ns | 99 | 86 | 0.910ns |
| SiSSM84958 | 41 | 48 | 0.550ns | 72 | 113 | 9.090* |
| SiSSM86906 | 37 | 49 | 1.670ns | 107 | 81 | 3.600ns |

<sup>a</sup>Chi-square = 3.841 at the 5% level ( $P < 0.05$ ).

**Figure 1.**
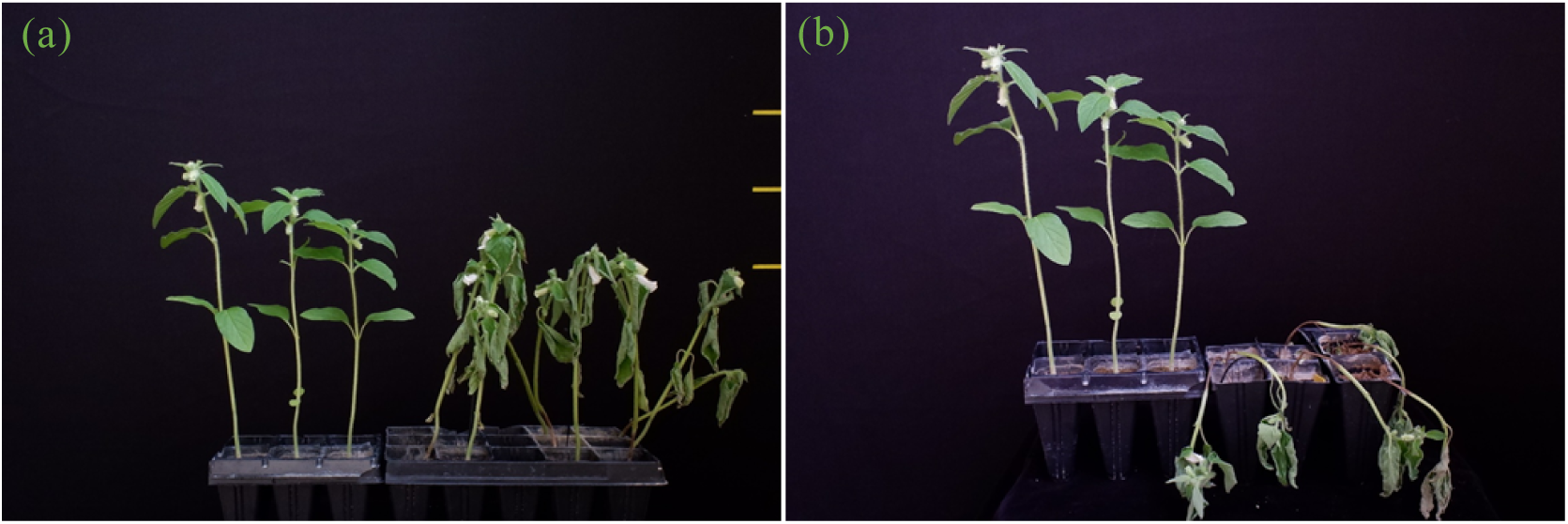
Plant wilting caused by *P. nicotianae*. (a) resistant and susceptible lines after seven days of inoculation (DPI) (b) resistant and susceptible lines after fourteen days of inoculation (DPI) with *P. nicotianae*.

**Figure 2.**
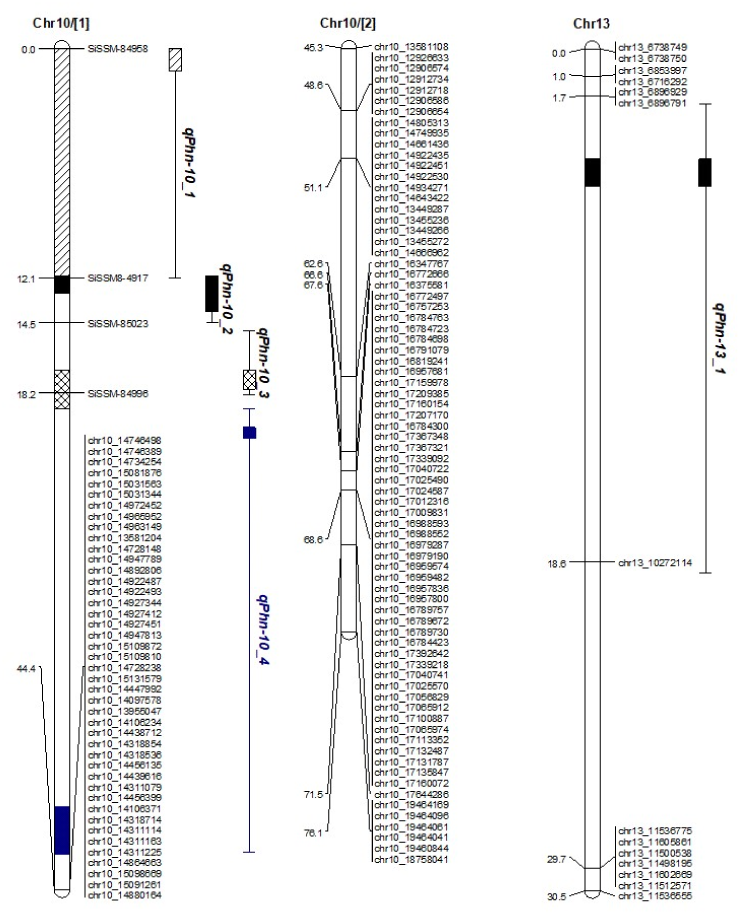
Linkage map of chromosome 10 showing significant QTLs for Phytophthora blight resistance in Goenbaek × Osan RIL population.

Besides the RIL populations, the germplasm lines (*n =* 87) were also evaluated for their pathogenic response to *P. nicotianae*. The infection range observed in the plants was categorized from 0–9 scale from *P. nicotianae* resistance to susceptible (Oh et al., 2018). Of the 87 lines, 21 lines were categorized as resistant (R), 4 lines as moderately resistant (MR), 12 lines as moderately susceptible (MS), and 50 lines as susceptible (S) to *P. nicotianae* isolate KACC48121. Similarly, the disease evaluation score for KACC48120 was 16 R, 1 MR, 7 MS, and 36 S, whereas those for No2040 were 16 R, 1 MR, 2 MS, and 47 S (Table S1).

### BSA analysis

In order to identify the markers linked to Phytophthora blight resistance, bulked segregant analysis (BSA) was performed in two segregating F_2_ populations. A total of 54 primer pairs (SSR) were used to detect polymorphism among the parental lines and resistant and susceptible bulks (Table S3). Out of 54 primers used, six primer pairs were polymorphic in two susceptible cvrs. Milsung and Osan and resistant cvr. Goenbaek parental lines. The similar polymorphic pattern showed in Population-I five primer pairs (SiSSM-84917, SiSSM-84958, SiSSM-84996, and SiSSM-85023 and SiSSM-86906), in Population-II six primer pairs (SiSSM-84917, SiSSM-84958, SiSSM-85023, SiSSM85354, SiSSM-85570 and SiSSM-86906) were polymorphic (Table 1; Table S3). In total seven SSR markers, SiSSM-84917, SiSSM-84958, SiSSM-84996, SiSSM-85023, SiSSM85354, SiSSM-85570 and SiSSM-86906 could discriminate between resistance and susceptible genotypes and their resistant and susceptible bulks in both the populations (Table S3). Among the surveyed primers in BSA analysis, six primers were able to distinguish parents and F_2_ bulks derived from the Goenbaek and Osan, Goenbaek and Milsung varieties (Figure S1) correlated with phenotypic disease evaluation scores. As a result, these primers were scored in order to know the correlation with the phenotypic disease evaluation scores for each individual of Population-I and Population-II (Table 1).

### Single marker analysis to identify markers associated with Phytophthora blight in Population-II

Single marker analysis was performed to identify markers associated with Phytophthora blight in Population-II. From the BSA, we identified six polymorphic SSR markers in which two SSRs, SiSSM-85354 (15,489,851 bp) and SiSSM-85570 (15,990,137 bp) showed polymorphism specific to Population-II (Figure S1). In this analysis, five markers (SiSSM-84917, SiSSM-85023, SiSSM-85354, SiSSM-84958, SiSSM-85570, and SiSSM-86906) (Table 1; Table S3) in Population-II were found to be linked to the Phytophthora blight in BSA analysis. Of these, SiSSM-84917 and SiSSM-85023 were strongly correlated with Phytophthora resistance (Table 2 and 3). The Kruskal-Wallis test analysis revealed that the probability of these SSRs to follow 1:1 binomial distribution and the markers flanking the QTL were also significant displayed through Single Marker Analysis (SMA) (P-<0.0001) (Table 2 and 3). Thus, this locus was temporarily designated *phn-10*.

**Table 2.** Kruskal-Wallis and Single marker analysis and validation of linked markers for Phytophthora blight resistance in 188 F_5:7_ RIL population derived from Goenbaek x Milsung.

| Marker ID | RR=R | rr=S | Physical position (bp) | cM | Kruskal-Wallis Analysis |  | Single Marker Analysis |  |  |
| --- | --- | --- | --- | --- | --- | --- | --- | --- | --- |
|  |  |  |  |  | K* | P-value | R<sup>2</sup>(%) | F-value | P-value |
| SiSSM84958 | 72 | 113 | 14635504-14663553 | 0.0 | 88.31 | <0.0001 | 25.31 | 0.11 | <0.0001 |
| *phn-10* | 89 | 98 | - | 4.3 | 186.0 | <0.0001 |  |  |  |
| SiSSM84917 | 90 | 94 | 14539434-14539477 | 6.6 | 130.60 | <0.0001 | 36.48 | 0.05 | <0.0001 |
| SiSSM85023 | 99 | 86 | 14765956-14765975 | 9.0 | 116.46 | <0.0001 | 32.00 | 0.08 | <0.0001 |
| SiSSM85354 | 89 | 95 | 15489851-15489880 | 12.1 | 105.01 | <0.0001 | 28.12 | 0.10 | <0.0001 |
| SiSSM85570 | 95 | 93 | 15990137-15990170 | 14.7 | 86.21 | <0.0001 | 23.21 | 0.13 | <0.0001 |
| SiSSM86906 | 107 | 81 | 18816333-18816378 | 20.0 | 1.80 | NS | (0.41)<sup>NS</sup> | 0.25 | 0.2019 |

**Table 3.** Kruskal-Wallis and Single marker analysis and validation of linked markers for Phytophthora blight resistance in 90 F_5:7_ RIL population derived from Goenbaek x Osan.

| Marker ID | RR=R | rr=S | Physical position (bp) | cM | Kruskal-Wallis Analysis |  | Single Marker Analysis |  |  |
| --- | --- | --- | --- | --- | --- | --- | --- | --- | --- |
|  |  |  |  |  | K* | P-value | R<sup>2</sup>(%) | F-value | P-value |
| SiSSM-84958 | 41 | 48 | 14635504-14663553 | 0.0 | 33.00 | <0.0001 | 12.09 | 0.12 | <0.0001 |
| SiSSM-84917 | 46 | 43 | 14539434-14539477 | 12.1 | 53.17 | <0.0001 | 15.75 | 0.07 | <0.0001 |
| SiSSM-85023 | 46 | 43 | 14765956-14765975 | 14.5 | 43.86 | <0.0001 | 12.50 | 0.11 | <0.0001 |
| SiSSM-84996 | 42 | 46 | 14728083-14728148 | 18.2 | 55.05 | <0.0001 | 17.48 | 0.05 | <0.0001 |

*P*-value: P-value greater than 0.05 was regarded as a non-significant (NS) association, whereas *P* value of less than 0.05 was considered as significant marker-trait association. % R2: R2 value represents the percentage of phenotypic variance for Phytophthora blight resistance explained by each significant SSR. <sup>NS</sup>-Unlinked SSR marker: SSR markers that were not found associated with Phytophthora blight in our BSA analysis.

### Genotyping-by-sequencing analysis of the RIL population and linkage map construction

A GBS was performed on 90 F_5:6_ RIL population derived from Goenbaek and Osan and data used for genetic mapping. Through GBS analysis, a total of 40.2 Gbp of DNA sequence reads were obtained. The average number and total length of raw reads for each sample were 4,228,098 and 0.43 Gbp, respectively. The demultiplexed sequences of the 96 samples were trimmed by eliminating the sequences of the barcode and adaptor, and removing the low-quality information. As a result, the average number and total length of trimmed reads were 3,039,597 and 285.7 Mbp, respectively, and 71.94% of the total raw data mapped to the reference genome *Sesamum indicum* (version 2.0) (http://ocri-genomics.org/Sinbase_v2.0). In the initial attempt, 1773 markers were selected to generate a linkage map for the sesame population. Nineteen of the markers remained unlinked (*i*.*e*., no recombinants observed) by applying the regression method in JoinMap 4.0 and therefore were removed. Meanwhile, in the initial map construction for the linkage analysis, 75 unlinked loci to any group that did not fit very well and showed suspect linkages with a recombination frequency larger than 0.6 in the calculation option of each group were not included in the map. A total of 1678 polymorphic loci (1674 SNPs and 4 SSRs) were scored in the RILs. The genetic map comprised thirteen LGs covering a total map length of 887.49 cM (Table S2) and the length of each chromosome ranged from 17.98 cM for the smallest linkage group of Chr7 to 114.03 cM for the largest one of Chr1. The average genetic distance between neighboring SNP markers was 4.69 cM; the largest gap between SNPs in each chromosome ranged from 8.22 cM in Chr7 to 32.25 cM in Chr9. The gaps were mostly caused by a lack of SNPs available to map between the two lines. (Table S2; Figure S2).

### Mapping of QTL for Phytophthora blight resistance

The composite interval mapping (CIM) analysis performed with 1674 informative SNPs and 4 SSRs identified eight QTLs (LOD > 2.0) for *P. nicotianae* (KACC48121). The SSR marker SiSSM-86906 was removed from the QTL analysis because this locus was found unlinked to any linkage group determined by JoinMap 4.0. The phenotypic variances (PVE%) explained by individual QTLs ranged from 2.293% to 69.24%. Four of these five QTLs were detected on Chr10. In the Population-I, a resistance locus was detected on Chr10 with the LOD range of 2.27 and 24.53 using CIM. The QTLs *qPhn-10_2* and *qPhn-10_3* were located between flanking markers SiSSM-84917-SiSSM-85023 and SiSSM-85023-SiSSM-84996, at nucleotide positions 14,728,083 bp and 14,539,434 bp, respectively, determined through sesame the updated genome assembly and annotation (http://ocri-genomics.org/Sinbase_v2.0). The additive effect of this locus was 0.85, 0.82, respectively, and the resistant allelic effect was from Goenbaek (Table 4). A single QTL *qPhn-13_1* detected on Chr13 showed a negative additive effect, meaning allele derived from susceptible parent ‘Osan’ (Table 4). The QTL *qPhn-10_2* showed the highest LOD score and explained 69.24% of total PVE. The physical positions of the QTLs detected in Goenbaek × Osan 90 F_5:7_ RILs were compared using the *S. indicum* ‘ver.2’ reference genome. Two genomic regions on Chr10 (14.54-14.77 Mbp) and 13 (6.74-10.27 Mbp) were associated with resistance to Phytophthora blight, and the QTLs located on Chr10 were considered the most significant. Linkage analyses revealed *Phn-10* was linked to polymorphic markers from the BSA F_2_ polymorphism survey (Figure S1). Goenbaek was the same resistance parent in the two RIL populations, the two loci were the same locus.

**Table 4.** Phytophthora blight resistance QTLs detected by composite interval mapping in Goenbaek x Osan 90 RIL population using WinQTL Cartographer.

| QTL name | Marker ID | Position (cM) | Physical position (bp) | LOD | PVE(%) | Additive |
| --- | --- | --- | --- | --- | --- | --- |
| *qPhn-10_1* | SiSSM-84958-SiSSM-84917 | 0.0 | 14635504 - 14539477 | 10.02 | 39.68 | 0.64 |
| *qPhn-10_2* | SiSSM-84917-SiSSM-85023 | 12.1 | 14539477 - 14765975 | 24.53 | 69.24 | 0.85 |
| *qPhn-10_3* | SiSSM-85023-SiSSM-84996 | 18.2 | 14765975 - 14728148 | 20.71 | 64.79 | 0.82 |
| *qPhn-10_4* | SiSSM-84996-chr10_14746498 | 42.2 | 14728148 - 14746498 | 2.27 | 16.78 | 0.42 |
| *qPhn-13_1* | Chr13_6896791-chr13_10272114 | 3.7 | 6896791-10272114 | 2.19 | 2.93 | -0.18 |

### Mapping SNPs and haplotype blocks of GWAS population

To validate the QTLs detected from the biparental populations, a GWAS study was performed using sesame accessions using the greenhouse-based layer test assay (Table S1). Using CMLM in the GAPIT R package PCA+K Bonferroni correction has captured associations for three *P. nicotianae* isolates with major effects on Chr10 (Table 6; Figure 4). The CMLM successfully detected significant markers using threshold FDR *P*-value < 0.001. A total of 44 SNPs on Chr10 were found to be associated with Phytophthora blight. Among 44 SNPs, 19 SNPs were associated and co-localized between *P. nicotianae* KACC48121, KACC48120, No2040 isolates, with coincided similar genomic region, with a range –log_10_(*P*) value 4.11 to 8.31, respectively. (Table 6; Figure 4). On Chr10, SNPs associated with *P. nicotianae* isolate KACC48121 detected between 14.56 and 15.10 Mbp corresponded to the location of the *qPhn-10_1, qPhn-10_2, qPhn-10_3, qPhn-10_4* QTLs (Tables 4 and 6; Figures 4 and 5). The data suggests the QTLs, *qPhn-10_1, qPhn-10_2, qPhn-10_3, qPhn-10_4* lies in between S10_14563939 and S10_15131579, which represents 0.0 to 44.4 cM. QTLs *qPhn-10_1, qPhn-10_2* detected by interval mapping (IM) in Population-II together with SIM (Tables 2 and 5), which represents 0.0 to 9.0 cM lies in between S10_14563939 and S10_15131579 (14.56-15.13 Mbp) detected by GWAS (Table 6). Overall, 14 SNP markers from GWAS associated with KACC48121 isolate were found to be overlapped with QTL *qPhn-10_4* in Population-I (0.44 cM) with PVE of 16.78% (Table 4 and 4; Figure 3). These 14 SNPs in genetic linkage map analysis grouped all together at 44.42 cM interval by JoinMap 4.0 (Figure 3). These 14 SNPs having similar distance (in cM) may be due to lower population size used for the current linkage map and narrow genetic base of two parental lines.

**Table 5.** Phytophthora blight resistance QTLs detected by interval mapping in Goenbaek x Milsung 188 RIL population using IM model in MapQTL6.

| QTL name | Marker ID | Position (cM) | Physical position (bp) | LOD | PVE(%) | Additive |
| --- | --- | --- | --- | --- | --- | --- |
| *qPhn-10_1* | SiSSM-84958 | 0.00 | 14635504 - 14663553 | 27.05 | 48.6 | -2.87 |
|  | SiSSM-84917 | 6.60 | 14539434 - 14539477 | 50.95 | 71.5 | -3.40 |
| *qPhn-10_2* | SiSSM-85023 | 9.00 | 14765956 - 14765975 | 40.57 | 63.2 | -3.20 |
| *qPhn-10_5* | SiSSM-85354 | 12.10 | 15489851 - 15489880 | 34.17 | 56.9 | -3.03 |
|  | SiSSM-85570 | 14.70 | 15990137 - 15990170 | 25.28 | 46.3 | -2.72 |

**Table 6.**
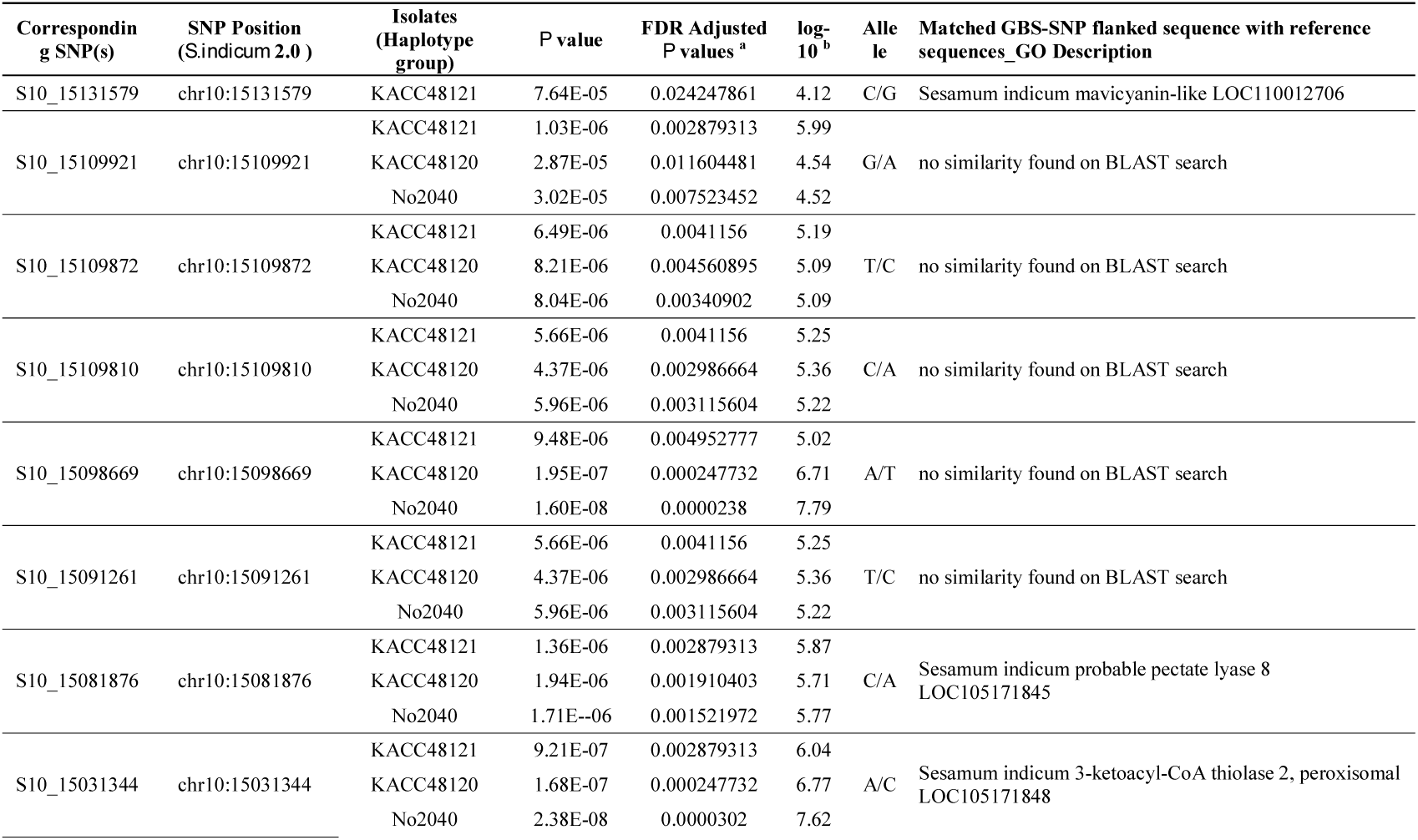
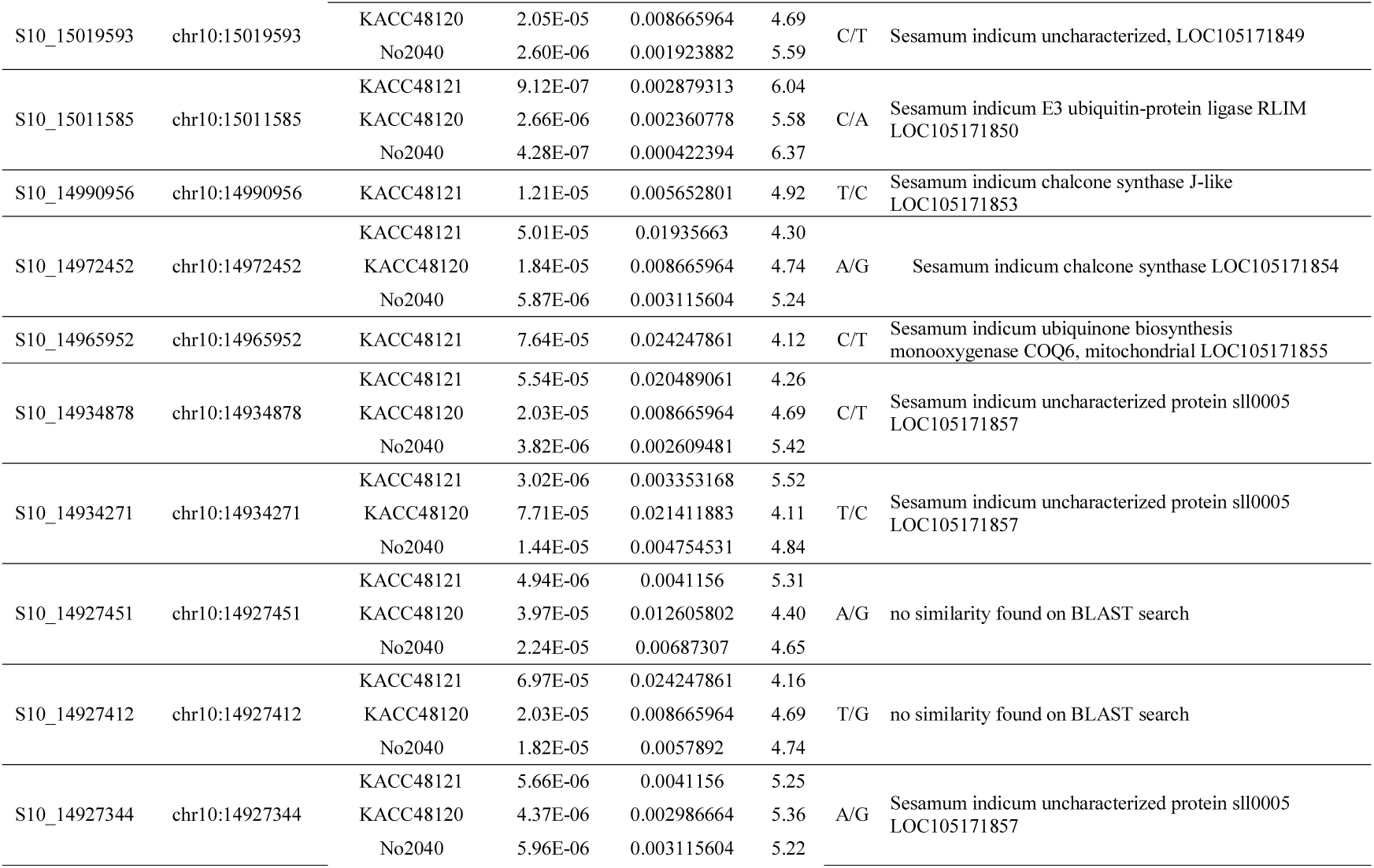
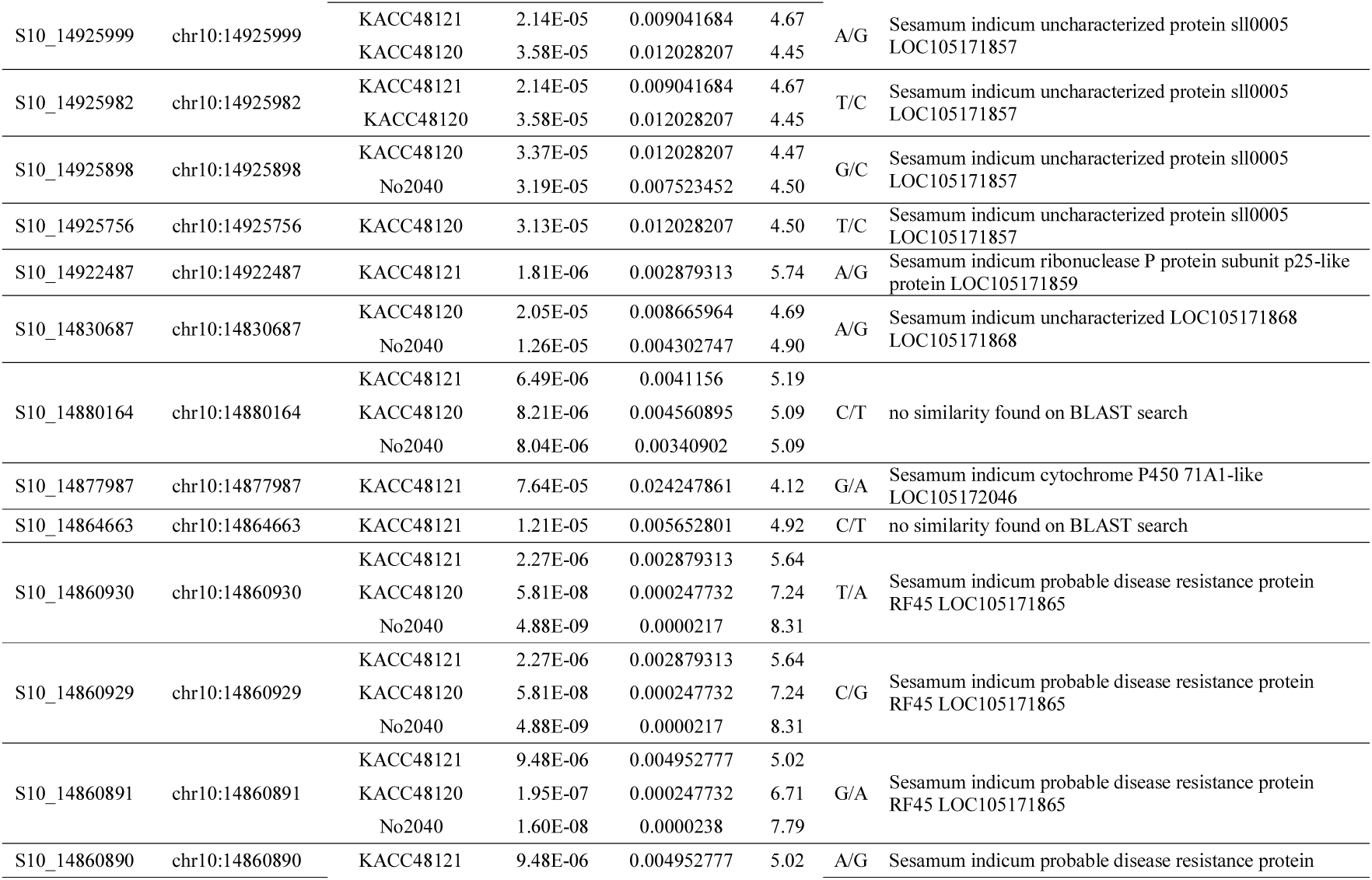
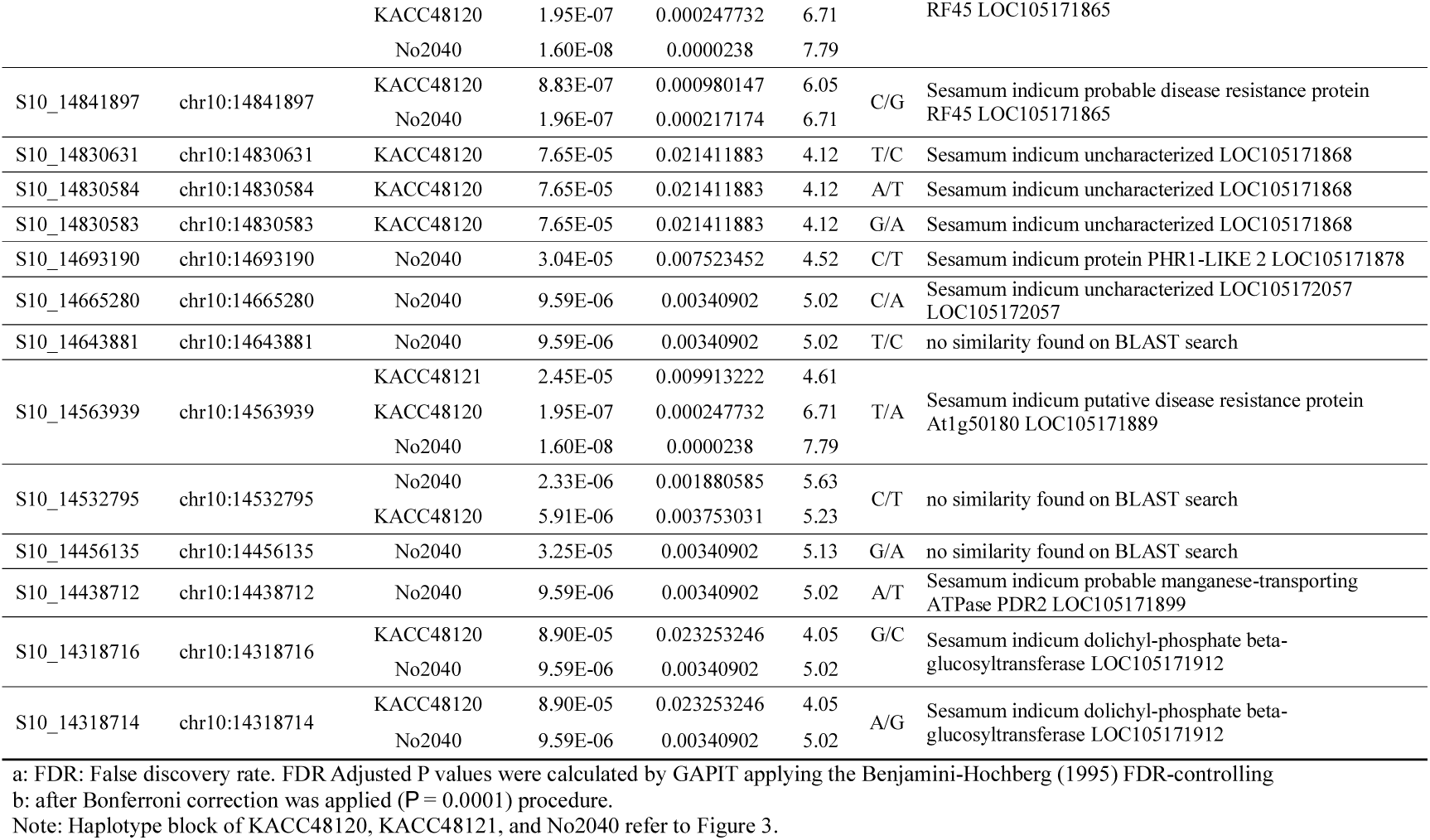
Candidate genes from the GWAS regions for *qPhn-10* and their Gene Ontology (GO) descriptions. LOC^§^ID given in the NCBI genome data base.

**Figure 3.**
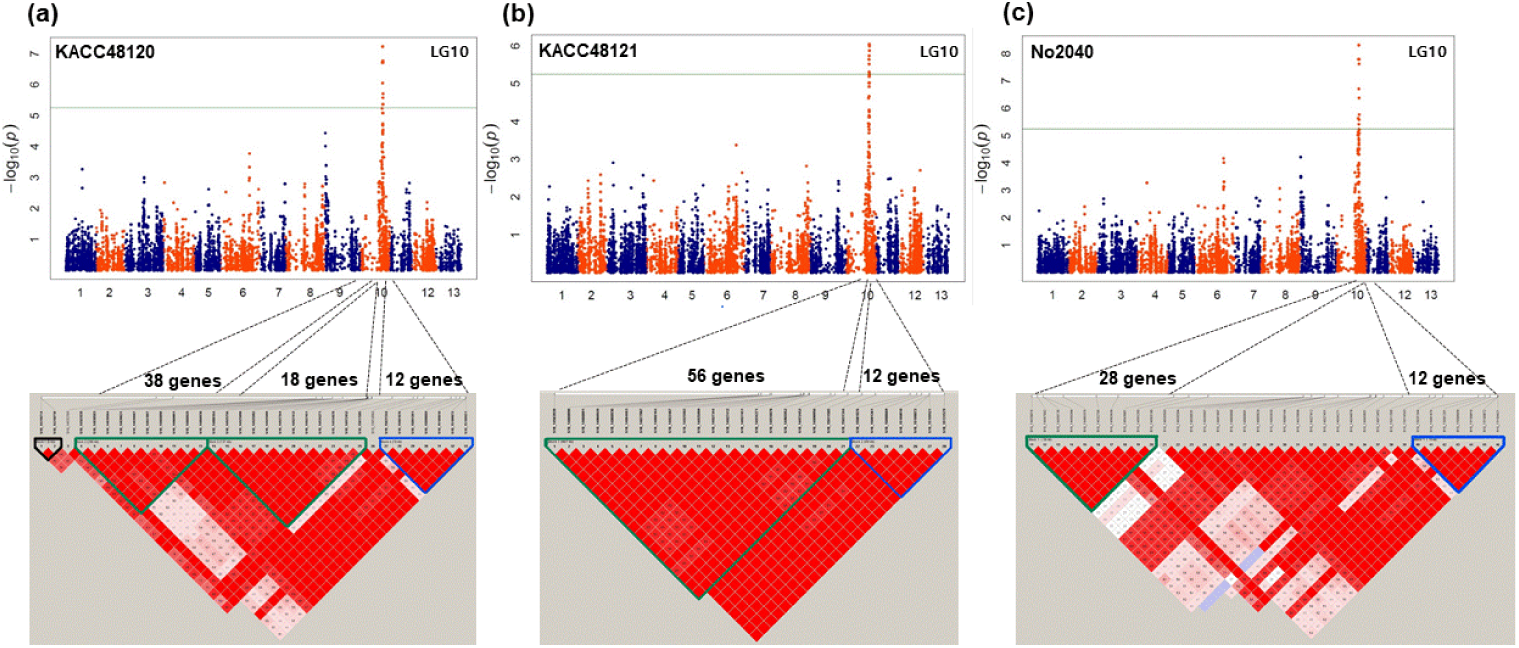
Manhattan plots and Haplotype block analysis based on GBS-GWAS showing the significant SNPs associated with *Phn* resistance. (a) Significant SNPs associated with isolate KACC48120. (b) Significant SNPs associated with isolate KACC48121 and (c) Significant SNPs associated isolate No2040 on Chr10.

**Figure 4.**
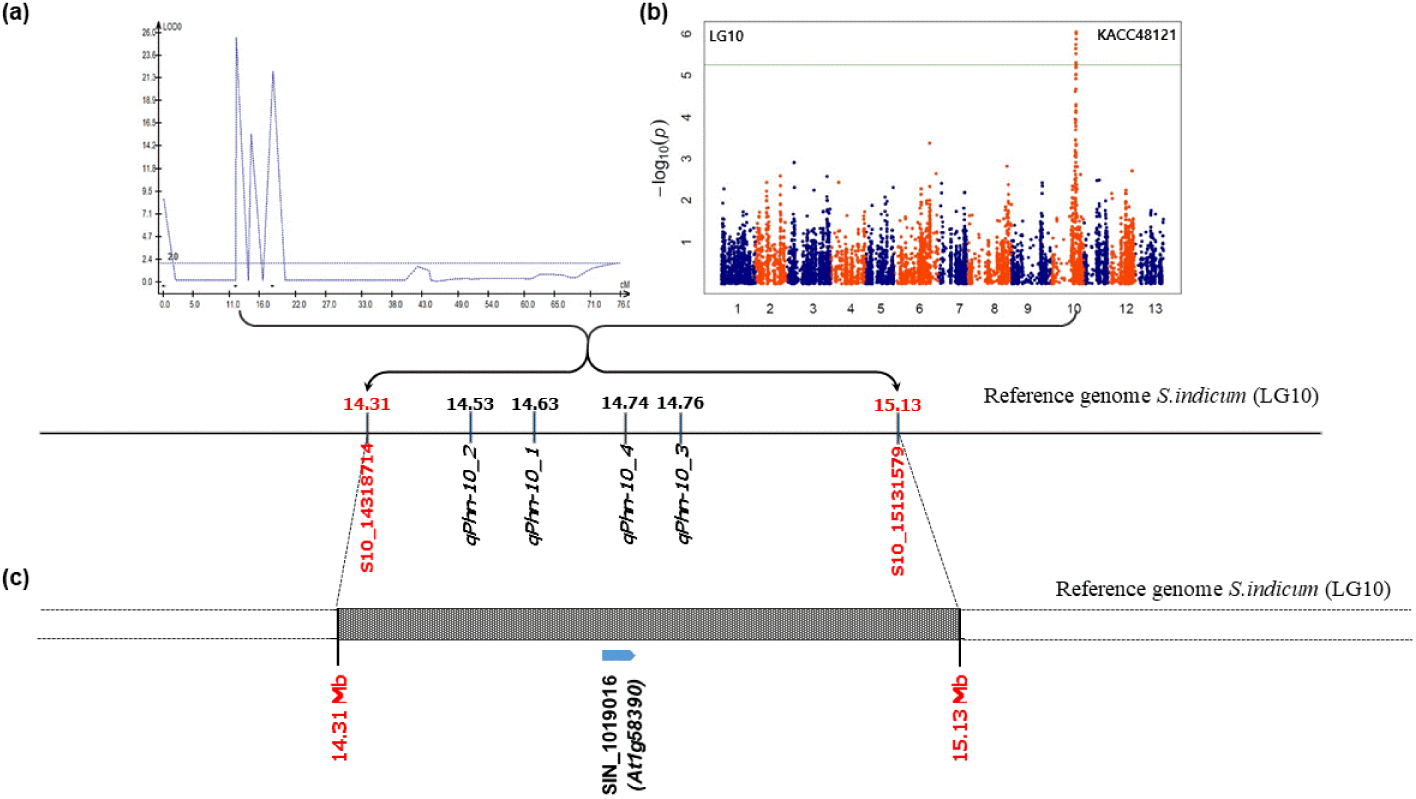
Identification of target region controlling Phytophthora disease resistance by linkage and association mapping. (a) QTLs associated with Phytophthora disease resistance in Population-I (Goenbaek ×Osan). (b) Manhattan plots of association analysis for Phytophthora disease resistance. Each dots represents SNP. The significant threshold –log^10^(*p*) = 8.31; (c) The target interval on Chr10. The blue bar represents the gene which was identified to exhibit a significant correlation between phenotype variation of Phytophthora disease resistance and gene expression levels.

**Figure 5.**
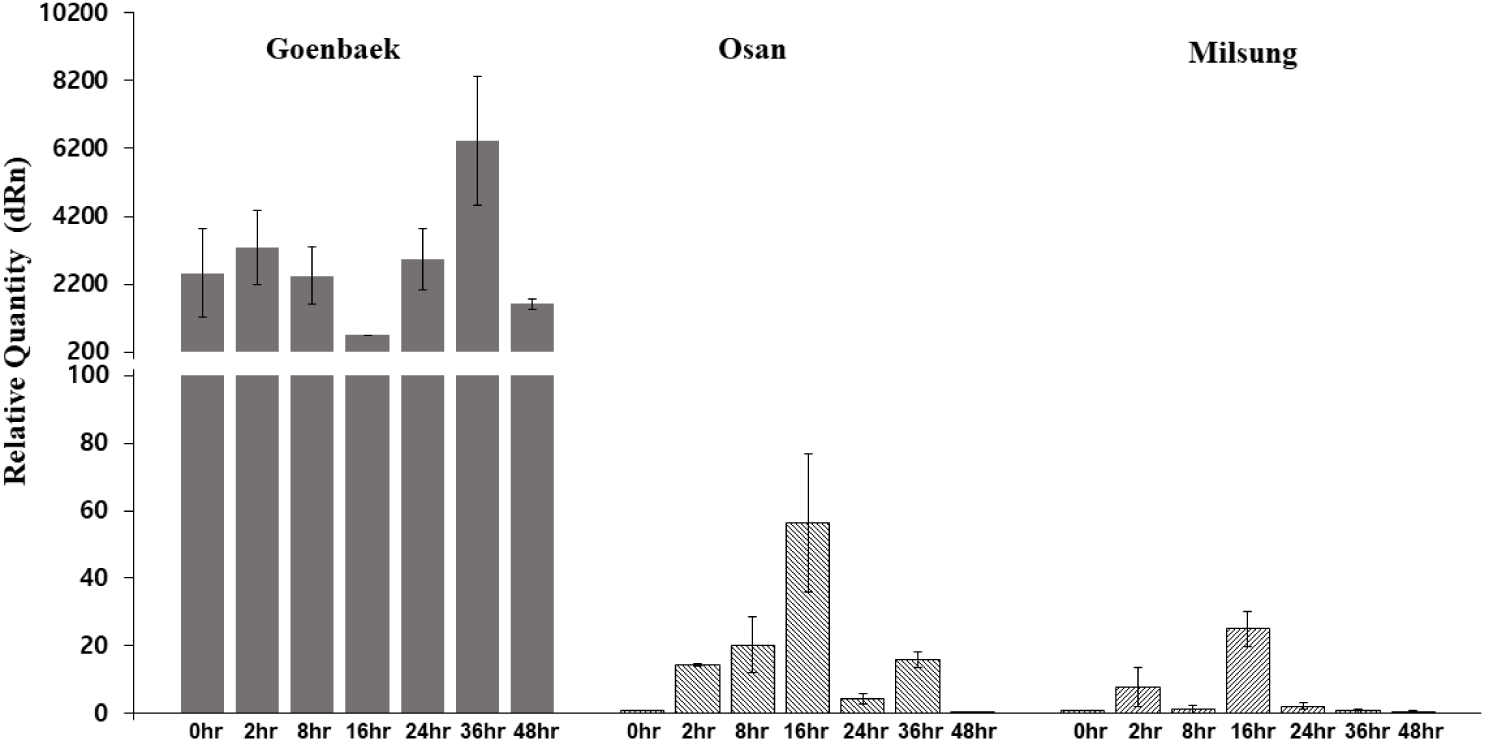
The quantitative real-time PCR (qRT-PCR) analyses of *SIN_1019016 (At1g58390)* and mRNA expression levels of *SIN_1019016* (*At1g58390)* probable disease resistance protein at seven time-points (0, 2, 8, 16, 24, 36, and 48) after inoculations for Goenbaek, Osan, Milsung sesame cultivars, respectively.

The raw data of flanking sequences (600 bp) obtained from GBS sequencing of associated SNPs were BLASTed for identifying candidate genes. The analysis of physical locations of associated SNP markers revealed that all markers were located in intergenic regions. Five SNPs (S10-14860929, S10-14860930, S10_14860891 and S10_14860890, S10_14841897) were associated with *P. nicotianae* isolates KACC48120, KACC48121 and No2040 were found to be in intron part within major resistance gene *SIN_1018978* and *SIN_1018979* (LOC105171865, probable disease resistance protein RF45) on sesame Chr10 using BLAST (Table S5). However, we noticed a single gene with 5 SNPs markers (S10_14860930, S10_14860929, S10_14860891, S10_14860890 and S10_14841897) found inside the gene as a candidate *R* gene and all are in intergenic region (Table 6). For a more in-depth evaluation of these regions of Chr10 containing markers, an analysis of haplotype block between the SNP markers in these intervals with candidate genes was carried out. Associated significant markers within this genomic region of haplotype block includes 6 and 7 SNP markers spanning 244.0 and 79 Kb for isolate No2040, 14, 13, and 7 SNP markers spanning 305, 136, and 79 Kb for isolate KACC48120, 27 and 7 SNP markers spanning 449 and 79 Kb for isolate KACC48121, respectively (Table S5; Figure 4 a,b,c). Haplotype block possessing marker(s) identified were significantly associated with a Phytophthora blight resistance.

We compared the linkage mapping and association mapping results. The QTL intervals *qPhn-10_1, qPhn-10_2*, and *qPhn-10_3* in Population-I (Table 5), QTLs *qPhn-10_1, qPhn-10_2* in Population-II were overlapped with GWAS QTL interval on Chr10 (Figure 5). These SSR markers not only exhibited the highest PVE value and had significant marker-trait correlation (*P* < 0.0001), but were also appeared in haplotype groups of No2040 (28 genes), KACC48120 (38 genes) and KACC48121 (56 genes) genomic region (Figure 4). We survey a number of candidate genes between SSR markers, SiSSM-84917 and SiSSM-85023 (14,539,477 bp to 14,765,975 bp) (Table S4). The genomic region between 44 associated SNP markers is 0.81 Mbp (14.31 – 15.13 Mbp) and is not large (Table 6). This QTL was considered major associated with four SSR markers mapped at 0.0 – 9.0 cM interval in a strong agreement with a physical distance of significant SNP markers associated with GWAS confirming tightly linked to disease resistance. Therefore, we focused on the genes in this interval. The corresponding genomic sequences of the region after haplotype block analysis were extracted. An annotation analysis showed that Chr10 from 14.31–15.13 Mb contains 80 genes (Table S5). These genes starting from the letter “*SIN*” were annotated according to the *S*.*indicum* v.2 reference annotation database (http://www.sesame-bioinfo.org/Sinbase2.0/).

### Identifying candidate genes associated with SSR markers

All the SNPs associated with Phytophthora blight were further evaluated by aligning the tag sequence having the associated SNP against all the known sequences to infer position and the potential candidate genes carrying a true causative polymorphism. Each SNP in intergenic position was considered for potential functional annotation based on the actual proximity of nearby located genes (Table S5; Table 6). The combined analysis involved BSA, linkage and association mapping approaches for identifying the possible *R* genes as strong candidates in this overlapped region (Table S4 and S5). Thirty-four putative genes were present in the corresponding genomic region. Of these, 9 genes encoded the NB-ARC domain (also identified as NBS-LRR proteins) found in many defense-related genes. Besides, 8 genes encoded protein kinase-like and Ser/Thr protein phosphatases, and 3 genes encoded pentatricopeptide (PPR) repeat-containing protein domains, both of which have been reported as stress-associated proteins.

The genes *SIN_1019026, SIN_1019021, SIN_1019020, SIN_1019018, SIN_1018958* encoding ubiquitin ligases and ubiquitin-related modifiers are shown to be involved in the regulation of pathogen-induced signaling since ubiquitination is known to play a critical role in the signaling pathways mediated by ABA (Stone, 2014). Five genes, *SIN_1019014, SIN_1018999, SIN_1018997, SIN_1018996, SIN_1018995* encoded F-box proteins, which have been reported to interact with plant disease hormones such as JA and promotes the expression of JA-responsive genes (Stotz et al., 2013). Remaining four genes known to (*SIN_1019019, SIN_1018986, SIN_1018975, SIN_1018974*) be a cytochrome P450 family protein gene were also reported plant disease resistance genes (Han et al., 2015). Of the identified candidate genes on Chr10 by haplotype block, we selected randomly five candidate genes, which were *SIN_1019016*, (probable disease resistance protein *At1g58390*; LOC110012696), *SIN_1019013* (proline-rich receptor-like protein kinase PERK3, LOC105171888), *SIN_1019001* (receptor-like protein kinase S.2, LOC105171879), *SIN_1018990* (putative disease resistance RPP13-like protein 1, LOC105172053) and *SIN_1018970* (vesicle-associated membrane protein 722, LOC105171860) for validation using qRT-PCR analysis to further confirm their role in Phytophthora resistance.

### qRT-PCR analysis

The functions of these genes were tested further using qRT-PCR analysis. The expression of the *SIN_1019016 (At1g58390)* gene in resistant line Goenbaek dramatically increased after 36 h of inoculation. It was interesting to note also at the 16 h of inoculation the resistant line Goenbaek was reduced expression whereas at the same time that inoculated in susceptible lines Milsung and Osan increased slightly, but at the 24, 36 and 48 h of post-inoculation showed reduced expression because susceptible plant tissue started wilting at that stage and eventually died (Figure 6). Goenbaek, Nuri, and RIL39 displayed almost no Phytophthora blight symptoms and had the highest expression from the susceptible lines (2 parental lines; Milsung and Osan, and 2 RILs; RIL26 and RIL34) (Figure S3). Reverse-transcriptase PCR analysis confirmed that the *SIN_1019016 (At1g58390)* gene had significantly higher expression in resistant line than in the Phytophthora non-resistant sesame cultivars and following RIL lines derived from Goenbaek (resistant) and Milsung (susceptible). Results of qRT-PCR and RT-PCR indicated the role of *SIN_1019016 (At1g58390)* gene Phytophthora blight disease resistance in sesame cultivars.

## Discussion

Molecular tagging of the gene(s) or QTL conferring Phytophthora blight resistance has not been previously reported in sesame. It is known that screening for disease resistance genes using molecular markers offers the additional advantage of permitting selection for resistance in the absence of the pathogen. Therefore, the availability of PCR-based markers will facilitate breeding efforts to efficiently develop Phytophthora blight-resistant sesame cultivars. In the present study, we employed an integrated approach involving QTL mapping using biparental mapping RIL populations, and GWAS to identify candidate genes governing Phytophthora blight resistance in sesame. In this study, using QTL mapping and single-marker analysis approaches, we identified a novel gene, designated as *phn-10* on Chr10. This QTL explained approximately 2.93–69.24% of the PVE, providing additional evidence of one minor and major gene coding for *P. nicotianae* resistance. The QTL was found to colocalize with the markers from BSA and with the QTL mapped using the Goenbaek × Osan (Population-I) and Goenbaek × Milsung (Population-II) populations. Several significant SSR markers associated with this major QTL found to cosegregates with Phytophthora blight resistance in the RILs used in this study. Thus, our study shows the success of GBS-generated SNP markers in gene-tagging in sesame.

From the GWAS results, a total of 44 SNPs were associated with Phytophthora blight resistance. Among 44 associated SNPs, 20 were common in all three isolates, 17 were co-localized in two isolate strains KACC48120 and No2040, three were common between two strains (KACC48121 and KACC48120) and 5, 5, and 3 were only specific to KACC48121, No2040, and KACC48120, respectively. It is possible that these common SNPs are highly correlated alleles conferring *R*-genes carrying plausible candidate genes could be referenced in QTL fine mapping of RILs with a favored combination for crossing with a hypersensitive parent to generate a fine-mapping population. However, associated SNPs to Phytophthora resistance for mining putative candidate disease resistance genes were located in intergenic regions not in exon regions. It is reported that the number of SNPs called from the GBS approach may not be large enough for GWAS and that SNP density in genic regions is much lower than that in intergenic regions (Ahn et al., 2018; Kim et al., 2014). Although, our germplasm panel subjected to GWAS identified sufficient and common SNPs associated with Phytophthora blight resistance to the three main strains (isolates KACC48121, KACC48120, No2040) which represents dominant types of strains in Korea and no resistance report in elsewhere before.

The QTL regions on Chr10, identified using biparental mapping as well as GWAS, were further scanned for identifying candidate genes within the interval. Thirty-four putative genes were found to be present in the corresponding genomic region. These genes encoded various domains, such as NBS-LRR proteins (having role in defense; (Meyers et al., 2003)), protein kinase-like and Ser/Thr protein phosphatases and PPR repeat-containing protein domains (stress-associated proteins), ubiquitin ligases and ubiquitin-related modifiers (regulation of pathogen-induced signaling; (Stone, 2014)), F-box proteins (interact with plant disease hormones such as JA and promotes the expression of JA-responsive genes (Stotz et al., 2013)), and cytochrome P450 family (plant disease resistance genes (Han et al., 2015)). These genes may have a role in imparting Phytophthora blight resistance in sesame. To further assess their role, we performed an expression analysis of *SIN_1019016* (*At1g58390*). The *SIN_1019016* gene was highly expressed in the resistant line Goenbaek compared to both the susceptible parent lines. Further, the expression of this gene found to dramatically increase after 36 h of inoculation (Figure 6), and reverted back to the normal levels at 48hr time point. At this time point, susceptible plant tissue eventually died. Thus, our qRT-PCR analysis showed that the upregulation of the *SIN_1019016* (*At1g58390)* gene in resistant line may have played a role in imparting Phytophthora non-resistance in the sesame cultivar Goenbaek. Expression analysis of other genes in the QTL intervals may provide additional molecular information regarding the role of this genomic region in controlling Phytophthora blight resistance in sesame.

NBS-LRR genes are reported to be the largest class of disease resistance (*R*) genes in plants, and are found as single (singletons) as well as clustered genes (tightly linked arrays of related genes) (Leister, 2004). Several R-genes encoding NBS-LRR proteins have been reported to cofer Phytophthora disease resistance in plants such as the *R1* gene for potato resistance to late blight potato (Ballvora et al., 2002), *Rps1* region containing Phytophthora resistance genes in soybean (Bhattacharyya et al., 2005), and the *Ph-3* gene from *Solanum pimpinellifolium* conferring resistance to *Phytophthora infestans* (Zhang et al., 2014). In this study, 9 genes from the QTL-associated genomic regions encoded NBS-LRR proteins, of which one gene was analyzed for its expression. This study is the first in Sesame showing role of *SIN_1019016* gene encoding NBS-LRR protein in conferring resistant to the Phytophthora races. Apart from *SIN_1019016* gene, other candidate genes from the QTL regions we identified in this study may also have significant role in Phytophthora blight disease resistance. Such genes when introgressed into the elite cultivars have been shown to impart resistance to respective pathogens. Therefore, precise nature, role, and organization of the *R*-genes have been a subject of research for long time. In-depth characterization of the genetic regions shown to be associated with the disease resistance to three races may reveal further information related to the molecular mechanisms involved in Phytophthora blight disease resistance in sesame.

## Conclusion

The present study showed that the use of GBS-based QTL identification in biparental mapping population coupled with GWAS provides means to identify sources of disease resistance in less-studied plant systems such as sesame. With this approach, we were able to identify SNPs/genomic regions significantly associated with Phytophthora blight resistance in sesame and confirmed by haplotype analysis One of the candidate genes from these regions showed higher expression in resistant parent indicating its role in Phytophthora blight resistance in sesame. Further molecular analysis of the QTL regions and candidate genes identified in this study could reveal insights into the molecular basis of Phytophthora blight resistance in sesame.

## Experimental procedures

### Plant materials

In this study, two RIL populations were developed by crossing a sesame cultivar Goenbaek as a resistant parent, with Osan (P2), and Milsung (P2), both being the susceptible parents. The cultivar Goenbaek was developed from a cross of Sungbun (mutated variety using gamma rays) and SIG950006-4-1-1 (crossing blocks) by the breeding team at the Rural Development Administration (RDA), National Institute of Crop Science Institute of Republic of Korea. In the year 2016, Goenbaek (P1) was crossed with Osan (P2), and Milsung (P2) were crossed at the Department of Southern Area Crop Science Breeding Station, National Institute of Crop Science (NICS), Miryang, Rep. of Korea to develop two different mapping populations. The subsequent F_1_ seeds were planted and self-pollinated in the greenhouse during the winter to produce F_2_ progeny. In 2017-18, the F_2_ seeds were planted in the greenhouse and self-pollinated to produce F_2:3_ family seeds, which were advanced to the F_5_ generation by single-seed descent. The F_5:6_ seeds were planted in the field of the National Institute of Crop Science, Miryang location in 2017 and were used as a source of bulk leaf tissue for DNA extraction and analysis. The F_5:5_ (Goenbaek×Osan; Population-I) and F_5:7_ (Goenbaek×Milsung, Population-II) mapping populations consisting of 90 and 188 lines along with the parental lines were germinated in sand-filled Stuewe Plug trays (Stuewe and Sons, Inc., Portland, Ore.), respectively.

For GWAS analysis, a set of 87 sesame germplasm lines, which also included Goenbaek, Osan, and Milsung, was utilized, and subjected for genotype-phenotype association mapping analysis (Table S1).

### Assessment of Phytophthora resistance in the germplasm panel and mapping population

Three *P. nicotianae* isolates (KACC48120, 48121, No2040) were selected from to test the seedling resistance of 87 sesame accessions, since they contained different virulence genes among each other and were the commonly used *P. nicotianae* isolates for resistance genetic analysis in Korea (Oh et al., 2018; Table S1). KACC48120, No2040, and KACC48121 were collected from Gyeongju (35°85’ N, 129°22’ E), Yecheon (36°65’ N, 128°45’ E) of Gyeongsangbuk-do provinces, and Naju, Jeollanam-do province (35°01’ N, 126°71’ E), respectively, in the sesame growing area of Jeollanam-do and Gyeongsangbuk-do provinces. The *P. nicotianae* isolate KACC48121 was used to analyze the genetic resistance of Phytophthora blight in our bi-parental population because it was virulent on more varieties than the other two isolates (Oh et al., 2018). Germinated seedling lines of 87 accessions, and the F_5:5_, F_5:7_ populations of Goenbaek x Osan, Goenbaek x Milsung placed in water-filled plug trays and each seedling were inoculated using five mL of zoospore (1×10^5^ zoospore /mL inoculum densities) suspension was soil-drenched at the basal stem of seedlings (V1 stage) using tube-connected bottle-top dispenser. The preparation of zoospore inoculum was the same as described previously (Oh et al., 2018). After 14 days, the inoculated types of each plant were recorded as resistant and susceptible. To resolve Phytophthora blight disease response into a Mendelian factor, “qualitative” mapping of phytophthora was conducted in the RIL population. Briefly, lines that had phytophthora blight indices showing leaf scars, wilting, stem damage and ultimate death were rated “resistant” or “0” and plants with indices of inoculation survivals were rated “susceptible” or “1”.

### DNA extraction and BSA analysis

In the present study, for disease rating, the symptom types were qualitatively categorized as either resistant (ratings of “0”) or susceptible (ratings of “1”). The difference between a resistant and susceptible response is the presence of chlorosis, necrosis of the roots and basal part of the stem, which indicates the successful reproduction of the pathogen conidia. The bulked segregant analysis (BSA) method was used for polymorphism screening (Michelmore et al., 1991), using equal quantities of DNA from 5 to 10 homozygous plants for each bulk. Two bulks of F_2_ plants were used for *P. nicotianae* mapping, including a *P. nicotianae* resistant bulk and a *P. nicotianae* susceptible bulk. Genomic DNA of the two parents, 90 F_5:6_ individuals, two (10 resistant and 10 susceptible) F_2_ bulks were extracted from young leaves using the CTAB methods as described by Paterson et al. (1999) with some modification to the components of the CTAB buffer (8.3 g NaCl, 2 g CTAB, 3 g PVP40, 5.92 g D-Sorbitol and 5Mm C_6_H_8_O_6_, 40mM C_5_H_11_NS_2_ in a total volume of 100 ml of 20 mM EDTA, 100 mM Tris-HCl, pH 8.0) to eliminate ultra-plentiful polysaccharides in sesame leaves. According to the initial mapping physical position, the sequence was downloaded from Sisatbase (http://www.sesame-bioinfo.org/SisatBase/), SSR markers in the targeted region were randomly selected and screened on the parental DNA samples. Initially, SSRs polymorphic between the bulks were used to screen the individuals within the F_2_ homozygous resistant (AA) class, and within the F_2_ homozygous susceptible (aa) class, and subsequently, the informative (i.e., parentally and BSA polymorphic) SSRs were used to screen the contrasting 90 F_5:5_ (Population-I) and 188 F_5:7_ (Population-II) individuals to determine if the marker of associated genomic region is linked for homozygous F_5:5_ of Population-I and F_5:7_ of Population-II plants for phytophthora resistance. A goodness-of-fit to the Mendelian segregation ratio was calculated using Chi-square (χ2) analysis to examine the segregation patterns of the phenotypes and selected SSR markers.

Chromosome-specific SSR markers were used to genotype individual F_2_, F_5:5,_ and F_5:7._ A 20 μl PCR reaction mixture consisted of 50 to 100 ng of sesame genomic DNA, 1X reaction buffer (25mM MgCl_2_ included), 100µM of each dNTPs, 10 pmol/µL forward and reverse primer, and 0.5 U of *e-Taq* DNA polymerase. All the chemicals used were purchased from SolGent Co., Ltd., Korea. The Chromosome-specific SSR marker PCR was performed in a DNA thermocycler (Mastercycler Nexus GSX 1 Eppendorf, Germany) using the following cycling parameters: 94°C (5 min); 35 cycles of 94°C (30 s), 55°C (30 s), 72°C (1 min); and final extension at 72°C (5 min). The PCR products were analyzed by 1.5% agarose gel electrophoresis and detected by DNA LoadingSTAR (DyneBio, Gyeonggi-do, South Korea). The primers were synthesized with Bioneer (Daejeon, Korea). The PCR products were also run on a QIAxcel capillary gel electrophoresis system according to the manufacturer’s instructions (Qiagen, Germany) and fragments were sized and scored using QIAxcel ScreenGel software (Qiagen, Germany).

### Preparation of libraries for genotyping-by-sequencing

The F_5:6_ and parents (resistant female parent Goenbaek and susceptible donor parent Osan) were genotyped with GBS technology using a protocol modified from Elshire et al. (2011) from SEEDERS Corporation (Daejon, South Korea). Using the varFilter command, SNPs were called only for variable positions with a minimum mapping quality (-Q) of 30. The minimum and maximum of read depths were set 3 and 100, respectively. In-house script considering biallelic loci was used to select a significant site in the called SNPs positions (Li et al., 2009). Depending on the ratio of SNP/InDel reads to mapped reads variant type are classified into three categories; homozygous SNP/InDel for more than 90%, heterozygous SNP/InDel for more than 40% and less than 60%, and the rest of them as etc. To control the quality of markers, missing proportion (MSP) <0.3 and minor allele frequency (MAF) > 0.05 were selected.

### Phenotype-genotype association analysis

The association panel used in this study (*n =* 87 sesame accessions) (Table S1) were subjected to GBS after filtering steps, a total of 8883 SNPs covering the whole genome with minor allele frequency >0.03 were used for GWA analysis. Before fitting the model, each marker was coded with the value 0 used for the reference allele and the value 1 for the alternative allele. Association analysis was performed for PCA and kinship matrices based on the Compressed Mixed Linear Model (CMLM). The CMLM was implemented using the R package Genomic Association and Prediction Integrated Tool GAPIT 2.0 (Lipka et al., 2012) for association mapping with 8883 SNPs in the germplasm panel of 87 accessions. Significance of probabilities generated in the association runs were transformed by -log_10_ *P* (FDR p value < 0.05). Scores for an individual chromosome were then inspected in Manhattan plots to determine whether the SNPs reached the significance threshold. After multiple test Bonferroni correction significance was defined at a uniform threshold of *p* < 5.54 × 10^−7^ (−log_10_ (*p*) > 6). The haplotype blocks and the identified significant SNPs for the chromosome were performed using the software PLINK v.1.9 and Haploview 4.2 (Barrett et al., 2005; Chang et al., 2015).

### Linkage map construction and candidate gene analysis

We selected the cross, Goenbaek×Osan, as the source of the segregating population to construct a genetic linkage map of sesame because of the sesame adequate frequency of SNPs. The genotype of each marker (SNP and SSR) and resistance gene locus for the 90 F_5:6_ RIL population (Population-I, Goenbaek × Osan) were used to construct a linkage map with JoinMap 4.0 (van Ooijen, 2006). The grouping mode was set as the independent limit of detection (LOD), the mapping algorithm was set as regression mapping, (recombination frequency < 0.4, LOD > 2.5 and jump =5), the map distances were calculated using Kosambi map function. Independence LOD and maximum likelihood algorithmwere used for grouping and ordering of markers respectively (van Ooijen, 2006). Missing data were considered a blank value to participate in the analysis. The linkage map was constructed using the SNP and SSR markers polymorphic between the two parents. The ordering of the markers within each chromosome was based on the recombination events between the markers, and the recombination distance was calculated using the Kosambi mapping function (Kosambi, 1943). Composite interval mapping (CIM) method was used to detect the QTLs, and the logarithm of odds (LOD) values greater than 2 was used to declare significant QTLs implemented in Windows QTL Cartographer 2.0 (Wang et al., 2011). A forward-selection backward-elimination stepwise regression procedure was used to identify co-factors for CIM. A 10-cM scan window was used for all analyses. The permutation tests were performed using 1,000 iterations to determine significant LOD thresholds. Furthermore, the additive effects and percentage variations explained (PVE) was obtained by performing CIM analysis. SNP, however, did not display frequent polymorphism; we continued to use it because of its frequent application in studies in humans and plants. The graphic representation of the linkage group and QTLs marked were created by Map Chart 2.2 (Voorrips, 2002).

Chromosome-specific marker data and the phenotypic disease evaluation scores were analyzed using interval mapping (IM) as implemented in MapQTL version 5.0 on Population-II (188 F_5:7_ RIL Goenbaek × Milsung). The genome-wide empirical thresholds for QTL detection (*P* < 0.05) were estimated using the permutation test as implemented in MapQTL version 5.0 (van Ooijen, 2004). The non-parametric Kruskal-Wallis test was also performed as a procedure of the MapQTL program version 5.0 (van Ooijen, 2004), in order to detect an association between the markers and traits individually.

The markers used in the mapping analysis were named based on the chromosome location and their physical region. For example, one “chr1_10766625” marker locus was named as “chr1” is chromosome 1 and “10766625” in the region on the physical map of sesame. Linkage groups and locus order were compared with published sesame linkage maps (Wang et al., 2015; Wang et al., 2016). QTL nomenclature was adapted from McCouch et al. (1997) in rice, which begins with “*q*”, followed by an abbreviation of the trait name, the name of the chromosome or linkage group, and then followed by the QTL number affecting the trait on the chromosome or linkage group (QTL + trait + number). For example, *qPhn*-10_1 designates the identified Phytophthora blight resistance QTL on chromosome 10. Hyphen “-” and the underscore “_” symbols stand before are for the chromosome in which the QTL is located and different QTLs for the same trait that are identified in one chromosome. Interactions among loci were not evaluated in this study since population sizes were too small to conduct meaningful tests.

### Kruskal-Wallis and Single Marker Analysis of variance (ANOVA)

We used single marker ANOVA and Kruskal-Wallis Analysis for validation of marker-trait association of detected QTLs. Briefly, for each of the SSR markers, the RILs were grouped according to the SSR, and then, one-way ANOVA was used to test for significant difference between the group means. *F*-statistics in the ANOVA and K* were used to define significant marker-QTL association. We used *P*-value <0.05 to define significant marker-QTL association. R-Square value from the ANOVA analysis was interpreted as a measurement of the proportion of phenotypic variation explained by the QTL.

### RNA extraction and qRT-PCR Analysis

Quantitative reverse transcription (qRT)-PCR was performed for the five most promising *R*-genes (Table S6) from linkage and association analysis described above. The gene-specific primers were designed based on the 5′-untranslated regions (UTR) and 3′-UTR of the gene models *SIN_1019016* (LOC110012696), *SIN_1019013* (LOC105171888), *SIN_1019001* (LOC105171879), *SIN_1018990* (LOC105172053), and *SIN_1018970* (LOC105171860) sesame reference genome *S_indicum*_v1.0. Total RNA was extracted from sesame plants at different time points after inoculation using RNeasy PowerPlant Kit (Qiagen) using the manufacturer’s instructions. Comparisons of gene expression in sesame samples were made based on a time series after sesame inoculation with *P. nicotianae* in two genotypes: Misung and Osan (susceptible) and Goenbaek (resistant) collected at 2, 8, 16, 24, 36 and 48 h post-inoculation. For each of these time-points the equivalent control (0 h), un-inoculated plant/tissue was also sampled for total RNA extract and qRT-PCR analysis. First-strand cDNA was synthesized from 1 μg of total RNA using Oligo(dT) RNA to cDNA EcoDry Premix (Takara, Dalian, China). Primers used for qRT-PCR were designed using Primer 3Plus online software (http://bioinformatics.nl/cgi-bin/primer3plus) based on their DNA sequences retrieved from Sesame reference genome database at the National Center for Biotechnology Information (NCBI). Primer Premier 5.0 software was used to assess the optimization of the primers. Finally, qRT-PCR reactions were conducted as described by the manufacturer for the RT system. Reaction mixtures (20 μl) contained the cDNA reverse transcription solution (1μl), PCR Master Mix (SYBR Green, Life Technologies LTD., Warrington, UK) and 0.2μM each of each primer.

The qRT-PCR thermocycling program was 50°C for 2 min, 95°C for 10 min, 40 cycles of 10s at 95°C, 10s at 60°C and 30s at 72°C. Then the melting curve temperature profile was obtained by heating to 95°C for 15s, cooling to 60°C for 1min, and slowly heating to 95°C for 15s with 0.5°C change of every 10s with continuous measurement of fluorescence at 520 nm. An 18S rRNA gene was used as an internal control to standardize the data, and the amount of target gene transcript was normalized compared to the constitutive abundance of sesame 18S rRNA (Noguchi et al., 2008). All reactions were performed in triplicate, including three non-template reaction as negative controls. Quantification of gene expression was performed using a 7900 Real-Time PCR System (Applied Biosystems, Foster City, CA, USA). Threshold values (CT) generated from the ABI PRISM 7900 Software Tool (Applied Biosystems, Foster City, CA, USA) were employed to quantify relative gene expression using the comparative 2^-ΔΔCT^ method (Livak and Schmittgen, 2001).

## Acknowledgments

This work was supported by a grant from the Rural Development Administration (Agenda project No. PJ01253601), National Institute of Crop Science (NICS), Republic of Korea.

## Author’s contributions

AS wrote the manuscript. OE conceived this project. KPK signed experiments, interpreted the results, critically evaluated and revised the manuscript. AS, KMS, KSU performed the experiments and analyzed the data. MHL, JIK, SBP, OKW, CKS helped to perform the experiments and collect the data. All authors read and approved the submission of this manuscript.

## Supporting information

Figure S1. Identification of SSR markers linked to *phn-10* using bulked segregant analysis. (a) SSR markers shown on resistance parent Goenbaek (PI), susceptible parent Milsung (P2), resistance bulk and susceptible bulk. (b) SSR markers shown on resistance parent Goenbaek (PI), susceptible parent Osan (P2), resistance bulk and susceptible bulk. Banding pattern of the resistance parent matches with the resistance bulk and the banding pattern of the susceptible parent matches with the susceptible bulk indicating that SSR markers are closely associated with *phn-10*.

Figure S2. Sesame linkage map constructed using 90 (F_5:7_) RILs derived from Goenbaek and Osan.

Figure S3. Expression of *SIN_1019016 (At1g58390)* gene in sesame un-inoculated lines. Detection of *SIN_1019016 (At1g58390)* gene in sesame lines, Goenbaek and Nuri were resistant cultivars, Milsung and Osan were susceptible cultivars, whereas, RIL26 and RIL34 were susceptible and RIL39 was resistant inbred lines of Population-II ((F_5:7_) Goenbaek and Milsung). 18rRNA was as actin was used as reference gene. To analyze each sample, 30 cycles were proceeded in RT-PCR.

Table S1. Details of the germplasm lines used in the association analysis.

Table S2. Segregation patterns and chi-square analysis in the F2 generation from the cross between Goenbaek and Osan (Population-I), Goenbaek and Milsung (Population-II) inoculated with KACC 48121.

Table S3. Detailed information of the genetic map developed using GBS-generated SNP markers.

Table S4. Details of SSR markers used for parental polymorphism survey in Population-I (Goenbaek×Osan) and Population-II (Goenbaek×Milsung).

Table S5. Genomic information and annotation information of the SNPs from SSR flanking regions associated with *phn-10* (*P*.*nicotianae*) resistance.

Table S6. The sequence matched with candidate genes using BLAST and *S. indicum* (v.2) reference genome for identification of putative genes within associated genomic region determined by GWAS and Haplotype analysis.

Table S7. List of oligonucleotides used for qRT-PCR analyses.

