## Supplemental Tables and Figures for "A Combinatorial Approach of Biparental QTL Mapping and Genome-Wide Association Analysis Identifies Candidate Genes for Phytophthora Blight Resistance in Sesame": Supplementary Figures_ A combinatorial approach of biparental qtl mapping and genome-wide association analysis identifies candidate genes for phytophthora blight resistance in sesame_Asekova S.docx

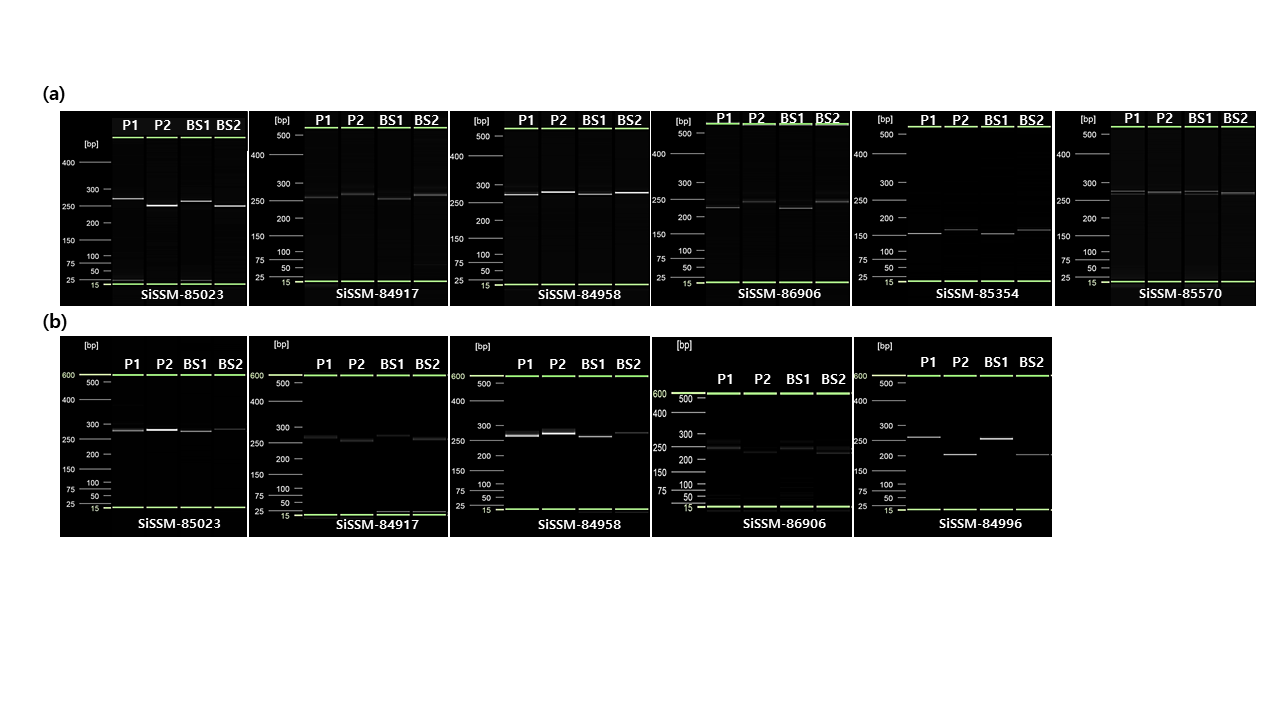

**Supplemental Figure S1.** Identification of SSR markers linked to *Phn-10* using bulked segregant analysis. (a) SSR markers shown on resistance parent Goenbaek (PI), susceptible parent Milsung (P2), resistance bulk and susceptible bulk. (b) SSR markers shown on resistance parent Goenbaek (PI), susceptible parent Osan (P2), resistance bulk and susceptible bulk. Banding pattern of the resistance parent matches with the resistance bulk and the banding pattern of the susceptible parent matches with the susceptible bulk indicating that SSR markers are closely associated with *Phn-10*.

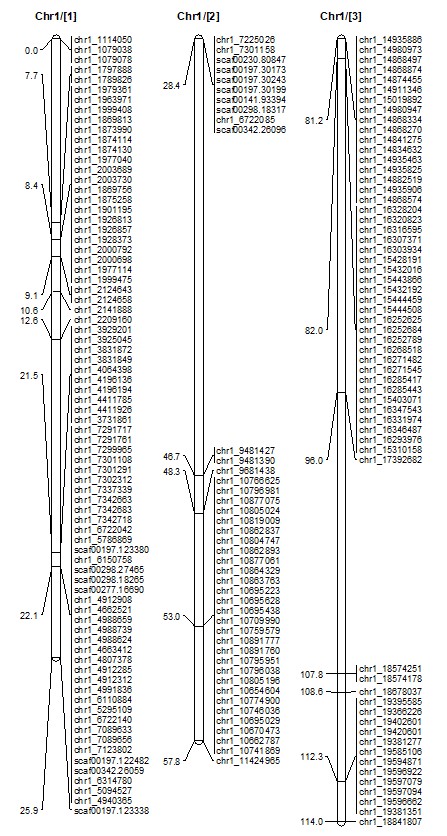

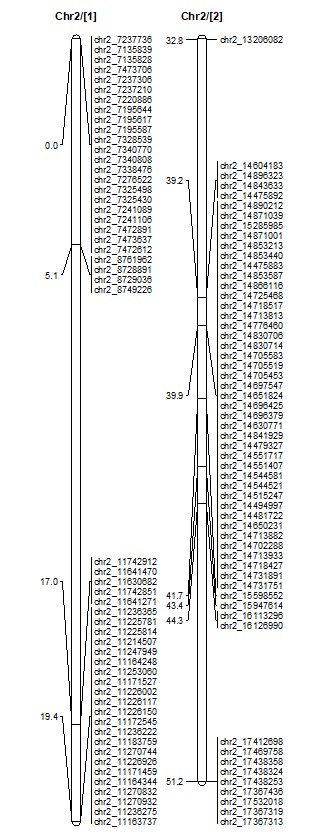

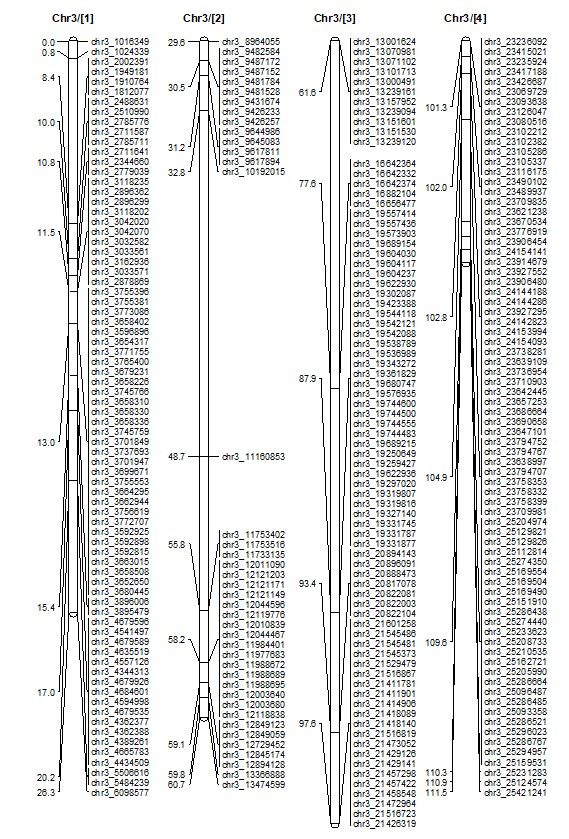

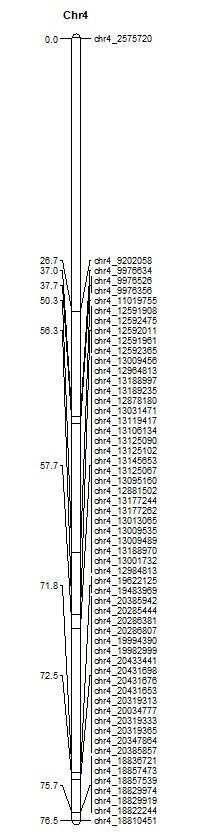

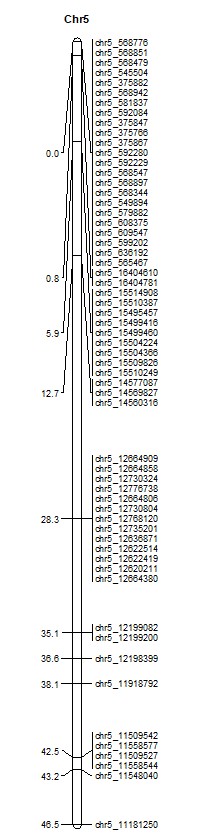

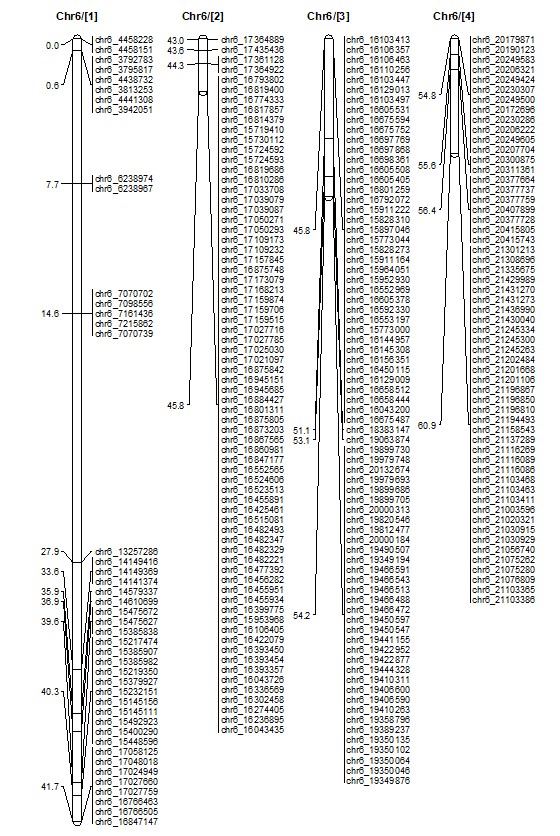

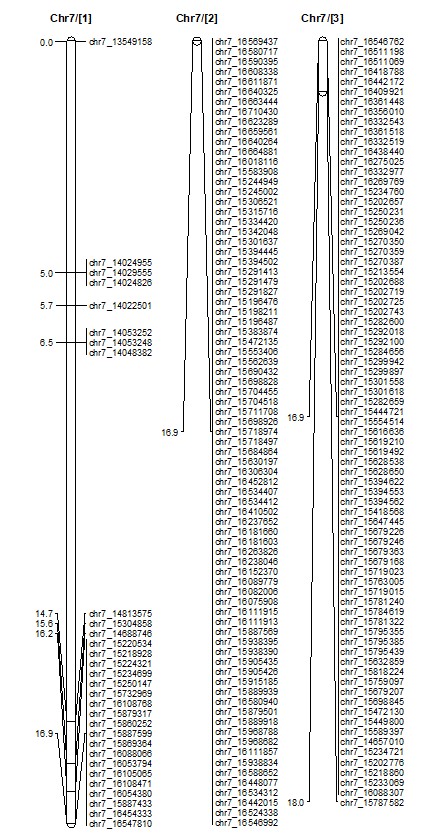

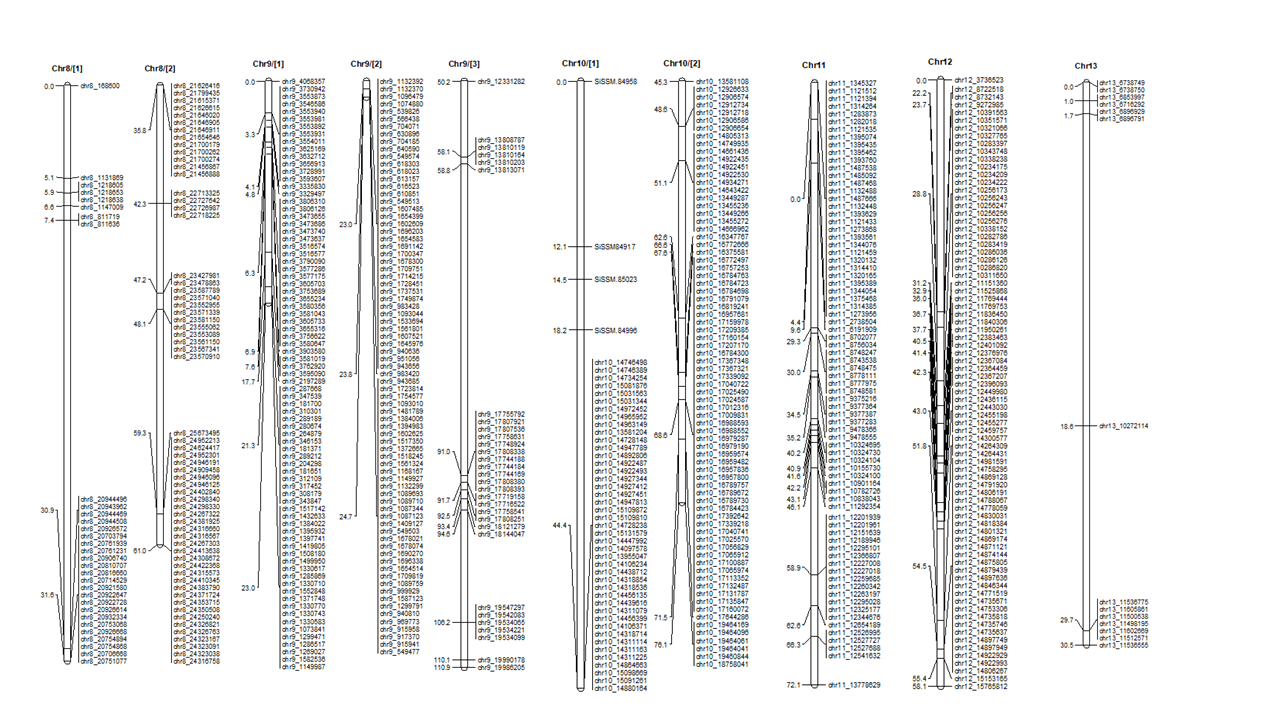

**Supplemental Figure S2.** Sesame linkage map constructed using 90 (F_5:7_) RILs derived from Goenbaek and Osan.

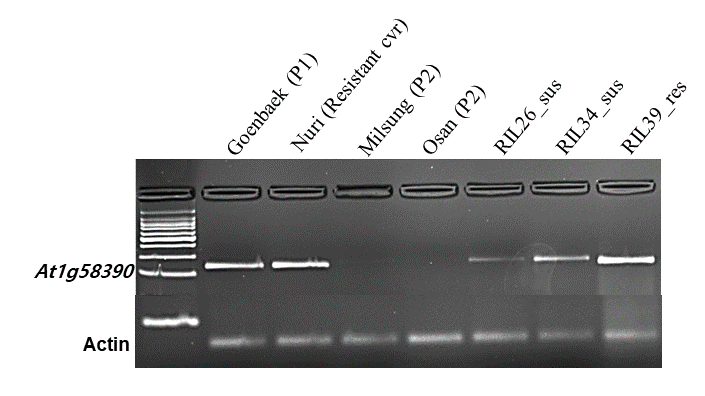

**Supplemental Figure S3**. Expression of *SIN_1019016* *(At1g58390)* gene in sesame un-inoculated lines. Detection of *SIN_1019016* *(At1g58390)* gene in sesame lines, Goenbaek and Nuri were resistant cultivars, Milsung and Osan were susceptible cultivars, whereas, RIL26 and RIL34 were susceptible and RIL39 was resistant inbred lines of Population-II ((F_5:7_) Goenbaek and Milsung). 18rRNA was as *actin* was used as reference gene. To analyze each sample, 30 cycles were proceeded in RT-PCR.
