## Supplemental Tables and Figures for "A Combinatorial Approach of Biparental QTL Mapping and Genome-Wide Association Analysis Identifies Candidate Genes for Phytophthora Blight Resistance in Sesame": Supplementary Tables_ A combinatorial approach of biparental qtl mapping and genome-wide association analysis identifies candidate genes for phytophthora blight resistance in sesame_Asekova S.docx

**Supplemental Table S1.** Details of the germplasm lines used in the association analysis.

| **Accession** | **Breeding status** | **Country of Origin** | **KACC48121** | | **KACC48120** | | **No2040** | |
| --- | --- | --- | --- | --- | --- | --- | --- | --- |
|  |  |  | **Disease score** | **Resistance level** | **Disease score** | **Resistance level** | **Disease score** | **Resistance level** |
| Dodam | Cultivar | Korea | 0.0 | R | - | - | 0.0 | R |
| Geonbaek | Cultivar | Korea | 0.0 | R | 0.0 | R | 0.9 | R |
| Superteagang | Commercial cultivar | Korea | 0.0 | R | - | - | - | - |
| Jinju | Cultivar | Korea | 0.0 | R | 0.0 | R | 0.1 | R |
| SIG960320-1A | Breeding line | Korea | 0.0 | R | - | - | 0.0 | R |
| Gochang collection | Landrace | Korea | 0.0 | R | - | - | - | - |
| Jinmi | Cultivar | Korea | 0.0 | R | - | - | - | - |
| Chamhwang | Cultivar | Korea | 0.0 | R | 0.0 | R | 4.5 | MR |
| Wild sesame | *S. alatum*/wild | Tropical Africa, India | 0.0 | R | - | - | 0.0 | R |
| Naman | Cultivar | Korea | 0.1 | R | 0.0 | R | 8.1 | S |
| Pungan | Cultivar | Korea | 0.4 | R | 0.2 | R | 1.3 | R |
| Namsan | Cultivar | Korea | 0.8 | R | 0.6 | R | 0.4 | R |
| Jungmo 5002 | Cultivar | Korea | 0.9 | R | 0.1 | R | 2.3 | R |
| Jinbaek | Cultivar | Korea | 1.4 | R | 0.0 | R | 0.4 | R |
| HS445-1-1-2-2-4 | Breeding line | Korea | 1.5 | R | - | - | 9.0 | S |
| Yoomi | Cultivar | Korea | 1.6 | R | 0.2 | R | 0.0 | R |
| Kangheuk | Cultivar | Korea | 1.8 | R | 0.9 | R | 7.2 | S |
| Sungbun | Cultivar | Korea | 1.8 | R | 7.9 | S | 9.0 | S |
| Kangheuk | Cultivar | Korea | 2.4 | R | 0.0 | R | 0.0 | R |
| Suwon | Cultivar | Korea | 2.6 | R | 8.9 | S | 9.0 | S |
| Hansan | Cultivar | Korea | 3.0 | R | 8.6 | S | 9.0 | S |
| Pungnyeong | Cultivar | Korea | 4.0 | MR | 8.1 | S | 9.0 | S |
| NonggiS-4 | Landrace | Japan | 4.3 | MR | - | - | 0.0 | R |

cont..

| **Accession** | **Breeding status** | **Country of Origin** | **KACC48121** | | **KACC48120** | | **No2040** | |
| --- | --- | --- | --- | --- | --- | --- | --- | --- |
|  |  |  | **Disease score** | **Resistance level** | **Disease score** | **Resistance level** | **Disease score** | **Resistance level** |
| Namda | Cultivar | Korea | 4.8 | MR | 8.1 | S | 9.0 | S |
| IT184749 | Landrace | United States | 5.0 | MR | 6.8 | MS | - | - |
| Pungseong | Cultivar | Korea | 5.2 | MS | 5.6 | MS | 9.0 | S |
| Yusung | Cultivar | Korea | 5.3 | MS | 7.9 | S | 9.0 | S |
| Seodun | Cultivar | Korea | 5.8 | MS | 8.8 | S | 9.0 | S |
| Ansan | Cultivar | Korea | 5.8 | MS | - | - | - | - |
| Yean | Cultivar | Korea | 6.0 | MS | 5.0 | MR | 9.0 | S |
| Heuksun | Cultivar | Korea | 6.2 | MS | 8.3 | S | 9.0 | S |
| Nambaek | Cultivar | Korea | 6.2 | MS | 8.6 | S | 9.0 | S |
| Hanseom | Cultivar | Korea | 6.2 | MS | 8.3 | S | 9.0 | S |
| Suzi | Cultivar | Korea | 6.3 | MS | - | - | 8.0 | S |
| Jinyul | Cultivar | Korea | 6.5 | MS | 5.3 | MS | 9.0 | S |
| Danbaek | Cultivar | Korea | 6.6 | MS | 8.1 | S | 9.0 | S |
| dt-45 | mutant | Israel | 7.0 | MS | 9.0 | S | 7.0 | MS |
| Pungsan | Cultivar | Korea | 7.4 | S | 8.3 | S | 9.0 | S |
| Yupoong | Cultivar | Korea | 7.4 | S | 0.1 | R | 0.0 | R |
| Manli | Cultivar | Korea | 7.6 | S | 8.2 | S | 9.0 | S |
| Mangeum | Cultivar | Korea | 7.8 | S | 7.2 | S | 9.0 | S |
| Yangan | Cultivar | Korea | 7.8 | S | 6.3 | MS | 9.0 | S |
| Jungmo 5003 | Cultivar | Korea | 8.0 | S | 0.4 | R | 0.9 | R |
| Manhuk | Cultivar | Korea | 8.0 | S | 8.6 | S | 9.0 | S |
| Hoeryong | Cultivar | Korea | 8.0 | S | 6.0 | MS | 9.0 | S |

cont…

| **Accession** | **Breeding status** | **Country of Origin** | **KACC48121** | | **KACC48120** | | **No2040** | |
| --- | --- | --- | --- | --- | --- | --- | --- | --- |
|  |  |  | **Disease score** | **Resistance level** | **Disease score** | **Resistance level** | **Disease score** | **Resistance level** |
| Dubeol | Cultivar | Korea | 8.1 | S | 0.0 | R | 9.0 | S |
| Baeksun | Cultivar | Korea | 8.2 | S | - | - | 9.0 | S |
| Jungmo 5007 | Cultivar | Korea | 8.4 | S | 8.3 | S | 9.0 | S |
| Gangan | Cultivar | Korea | 8.6 | S | 7.0 | MS | 7.9 | S |
| Yubaek | Cultivar | Korea | 8.6 | S | 6.4 | MS | 9.0 | S |
| Sangbaek | Cultivar | Korea | 8.6 | S | 0.0 | R | 0.0 | R |
| Gwangsan | Cultivar | Korea | 8.6 | S | 8.8 | S | 9.0 | S |
| Milsung | Cultivar | Korea | 8.6 | S | 7.2 | S | 9.0 | S |
| Galmi | Cultivar | Korea | 8.6 | S | 8.9 | S | 9.0 | S |
| Daheuk | Cultivar | Korea | 8.8 | S | 8.9 | S | 9.0 | S |
| Soonheuk | Cultivar | Korea | 8.8 | S | 7.6 | S | 9.0 | S |
| Hwangbaek | Cultivar | Korea | 8.8 | S | 8.8 | S | 9.0 | S |
| Pungnam | Cultivar | Korea | 8.8 | S | 8.7 | S | 9.0 | S |
| Hogoen | Cultivar | Korea | 8.8 | S | 7.6 | S | 9.0 | S |
| Anbaek | Cultivar | Korea | 8.8 | S | 7.6 | S | 9.0 | S |
| Yangheuk | Cultivar | Korea | 8.8 | S | 8.4 | S | 9.0 | S |
| PI157156 | Landrace | India | 9.0 | S | - | - | - | - |
| dt-sel | mutant | Israel | 9.0 | S | 9.0 | S | 6.8 | MS |
| Gomazou | Cultivar | Japan | 9.0 | S | - | - | 9.0 | S |
| Kyeonbuk 27 | Breeding line | Korea | 9.0 | S | - | - | - | - |
| Olleh | Commercial cultivar | Korea | 9.0 | S | - | - | - | - |
| Miho | Cultivar | Korea | 9.0 | S | - | - | - | - |

cont…

| **Accession** | **Breeding status** | | **Country of Origin** | **KACC48121** | | **KACC48120** | | **No2040** | |
| --- | --- | --- | --- | --- | --- | --- | --- | --- | --- |
|  |  |  |  | **Disease score** | **Resistance level** | **Disease score** | **Resistance level** | **Disease score** | **Resistance level** |
| Maniheuk | Commercial cultivar | Korea | | 9.0 | S | - | - | - | - |
| Superansan | Commercial cultivar | Korea | | 9.0 | S | - | - | - | - |
| Annam | Cultivar | Korea | | 9.0 | S | 8.4 | S | 9.0 | S |
| Buloheuk | Cultivar | Korea | | 9.0 | S | - | - | - | - |
| Heukjanggun | Commercial cultivar | Korea | | 9.0 | S | - | - | - | - |
| Gopum | Cultivar | Korea | | 9.0 | S | 8.7 | S | 8.0 | S |
| Baengmi | Commercial cultivar | Korea | | 9.0 | S | - | - | - | - |
| PI490033 | Landrace | Korea | | 9.0 | S | - | - | - | - |
| Plusansan | Commercial cultivar | Korea | | 9.0 | S | - | - | - | - |
| Kangbaek | Cultivar | Korea | | 9.0 | S | 0.1 | R | 0.8 | R |
| Osan | Cultivar | Korea | | 9.0 | S | 8.5 | S | - | - |
| Kyeonbuk 33 | Breeding line | Korea | | 9.0 | S | - | - | - | - |
| Miheuck | Cultivar | Korea | | 9.0 | S | 8.0 | S | 9.0 | S |
| Yunheuk | Cultivar | Korea | | 9.0 | S | 9.0 | S | 9.0 | S |
| Hwanggeum | Cultivar | Korea | | 9.0 | S | - | - | - | - |
| Arum | Cultivar | Korea | | 9.0 | S | 8.2 | S | 9.0 | S |
| Pyoungan | Cultivar | Korea | | 9.0 | S | 8.0 | S | 9.0 | S |
| PI279536 | Landrace | Mexico | | 9.0 | S | - | - | - | - |
| PI599446 | Landrace | United States | | 9.0 | S | 9.0 | S | 9.0 | S |
| Early Russian | Cultivar | United States, Texas | | 9.0 | S | - | - | - | - |

**Supplemental Table S2.** Segregation patterns and chi-square analysis in the F_2_ generation from the cross between Goenbaek and Osan (Population-I), Goenbaek and Milsung (Population-II) inoculated with KACC 48121.

| Cross (Generation) |  | Resistant (R) |  | Susceptible (S) |  | Total |  | Segregation ratio | χ^2^ |  | *P* |
| --- | --- | --- | --- | --- | --- | --- | --- | --- | --- | --- | --- |
| Goenbaek/Osan (F_2_) |  | 128 |  | 333 |  | 461 |  | 1:3 | 1.88 |  | 0.18 |
| Goenbaek/Milsung (F_2_) |  | 92 |  | 327 |  | 419 |  | 1:3 | 2.07 |  | 0.16 |

Note: df=1.0; χ^2^(0.05, 1) =3.84

**Supplemental Table S3.** Detailed information of the genetic map developed using GBS-generated SNP markers.

| **Linkage groups (LGs)** | **Number of markers** | **Map length**  **(cM)** | **Average marker distance**  **(cM)** |
| --- | --- | --- | --- |
| LG1 | 179 | 114.03 | 5.70 |
| LG2 | 110 | 51.15 | 5.12 |
| LG3 | 271 | 111.54 | 3.38 |
| LG4 | 58 | 76.53 | 7.08 |
| LG5 | 60 | 46.54 | 4.65 |
| LG6 | 245 | 60.94 | 2.90 |
| LG7 | 186 | 17.98 | 2.25 |
| LG8 | 92 | 61.03 | 5.09 |
| LG9 | 186 | 110.94 | 5.04 |
| LG10 | 122 | 76.12 | 5.44 |
| LG11 | 76 | 72.14 | 4.51 |
| LG12 | 79 | 58.09 | 3.63 |
| LG13 | 14 | 30.46 | 6.09 |
| Total | 1678 | 887.49 | 4.69 |

**Supplemental Table S4.** Details of SSR markers used for parental polymorphism survey in Population-I (GoenbaekⅹOsan) and Population-II (GoenbaekⅹMilsung)

| **Marker ID** | **Source** | **SSR motif** | **Size** | **Primer Forward (5'-3')** | **Primer Reverse (5'-3')** | **Start (bp)** | **End (bp)** |
| --- | --- | --- | --- | --- | --- | --- | --- |
| SiSSM79649 | SisatBase | (AG)19 | 189 | TTGTTTTGGTCGGTGTGTGT | CAAGGTTGATGTGGATGGTG | 253057 | 253094 |
| SiSSM79880 | SisatBase | (TA)14(GA)10 | 108 | AGAATTCGCATAGGTGTGGG | CTCGTATTATCCGAAACCGC | 702935 | 702982 |
| SiSSM80459 | SisatBase | (GA)17 | 280 | GCCTGGAAACGGTTTTAGATT | TCCAAACCCTAACGAAAGAGA | 1869257 | 1869290 |
| SiSSM80534 | SisatBase | (AAT)16 | 256 | TGCAGATCATTAAACCTTGAAAAA | TGGTCCAAATCTTTACGAGATG | 2007111 | 2007158 |
| SiSSM80636 | SisatBase | (AG)22 | 150 | GTCAATTCCCTTGTCCGAAA | AAGATTAGATGCGCCCTCAA | 2187273 | 2187316 |
| SiSSM81107 | SisatBase | (TA)8(GA)13 | 238 | CCATTGAATTTGATTGAGTCTAAGG | GGCCTTTTCTTCCGTCTAGC | 3127756 | 3127797 |
| 1643 | PMDBase | (AAT)8 | 273 | AAGGTTCGTCGCGTGAATAC | AAAGAGATCACGTGTCAAGCA | 3184346 | 3184369 |
| SiSSM81406 | SisatBase | (AT)15 | 240 | TCATTTCTAATCCAAAGGCTGAA | ACCCATATCCCCTATGGAGC | 3717471 | 3717500 |
| SiSSM81434 | SisatBase | (GA)15 | 267 | AGCACCAAATGCCAACTACC | ACCTGCGTGAGGAGAGAAGA | 3755082 | 3755111 |
| 2040 | PMDBase | (AG)14 | 272 | TTATTTGCCGATAGAGACGAA | AGGTGATTGTTTTCCTGCCA | 3955904 | 3955931 |
| SiSSM81764 | SisatBase | (AT)17 | 240 | AATTGAACACAAGCCCAAGG | TGCCAGCAAGAACGATTATG | 4370928 | 4370961 |
| 2326 | PMDBase | (AG)15 | 191 | TGAGAGAGGGTGATTTTGGG | ACAACCTGACCCAACCACAT | 4550913 | 4550942 |
| SiSSM82104 | SisatBase | (TA)11(GA)11 | 200 | GAGGCCAAAGCTGTTTTCAG | ACATTGTGACGAACTGCTGC | 5364697 | 5364740 |
| 2705 | PMDBase | (AT)18 | 251 | TTGAGTGTGCTTCGTTGGTT | GAGGCAGAAGCTGTTTCACC | 5768317 | 5768352 |
| 2907 | PMDBase | (GA)7gg(GA)9 | 270 | CAAAATTGCAGAAATTCCCC | TTCTGCCTTTTGTTTTCTGTGA | 6566921 | 6566954 |
| 3184 | PMDBase | (AAT)10 | 271 | GGAGGGAAAGGAATTTGAGG | GGGCAACAAGAAAATGGAGA | 7915210 | 7915239 |
| 3627 | PMDBase | (AT)7aataagaaag(GA)9 | 239 | GCATAAAGGGGCTAAATGAAA | CCCCACCTTCCACACTTAAC | 9886626 | 9886667 |
| 3722 | PMDBase | (AAT)11 | 181 | TTATGTCAAAATTTAGTGATTCCTATG | TCGGGTGCTCGAATTTTAAG | 10192859 | 10192891 |

cont…

| **Marker ID** | **Source** | **SSR motif** | **Size** | **Primer Forward (5'-3')** | **Primer Reverse (5'-3')** | **Start (bp)** | **End (bp)** |
| --- | --- | --- | --- | --- | --- | --- | --- |
| 3829 | PMDBase | (AT)14 | 150 | TTGCCTCATCATAAGCACAAA | GAACTGCAGGTTTCAGAGGG | 10672647 | 10672674 |
| 4014 | PMDBase | (AT)17 | 227 | TGAACCAAAAAGCATATTGATAGAA | AAATGTTCACGACATGCACC | 11284786 | 11284819 |
| SiSSM83576 | SisatBase | (ATT)14 | 261 | GTTCGGAAATATGCCCAAGA | AAGAGGAGCGTGAGCATTTC | 11439728 | 11439769 |
| 5108 | PMDBase | (AT)18 | 280 | GGTTGTGCACTAAGGAGTTGC | CAGCTGCACCAACAAAAGTG | 13847192 | 13847227 |
| SiSSM84895 | SisatBase | (AT)12 | 214 | TTCTTGGTTGAACTTTGTCTGG | GAATGAATGGATGCAGCAAG | 14513502 | 14513535 |
| **SiSSM84917^§^** | **SisatBase** | **(AG)22** | **253** | **GCACACATACACACACGCAC** | **TGTACCACATTCGCCTTGTC** | **14539434** | **14539477** |
| SiSSM84922 | SisatBase | (TA)17 | 219 | TTTCATGAAGCAGCAGAGGA | CGCGATGCAATTTATAACCC | 14570408 | 14570441 |
| SiSSM84923 | SisatBase | (A)12 | 260 | TGTTGATCTCTACCCACAACAAA | GATGACGAGACGAGTTTTTCAA | 14576365 | 14576376 |
| SiSSM84926 | SisatBase | (AG)8 | 241 | GCAAAGCAAGAATAGCAGGG | GGAAAGAAGAGCCCCAAAAC | 14586922 | 14586937 |
| SiSSM84927 | SisatBase | (A)10 | 276 | ACGCCCGCTCTTAACTACG | TCTTCTTCCCCCTCCTCTTC | 14588044 | 14588053 |
| SiSSM84930 | SisatBase | compound | 178 | AGTGGCGATCCCAATTTTAG | TTTATGGGTCCAAATCGACG | 14596288 | 14596393 |
| SiSSM84933 | SisatBase | (TG)7 | 239 | GCTTAGGCAAGACCATGTGC | TTCCAGTGGATTCATAGCCC | 14598782 | 14598795 |
| SiSSM84939 | SisatBase | compound | 268 | GAACCGGTGAAAACTTGTCC | ACGACTCTTCCACCCACAAA | 14605708 | 14605789 |
| **SiSSM84958** | **SisatBase** | **(CT)15** | **166** | **AGACCCTTTTAGGCTCTGGC** | **AAGCGCTGGATGGAATAGAA** | **14635504** | **14635533** |
| SiSSM84987 | SisatBase | (AC)7 | 259 | GGGTTTGATCCCATGTTGAC | ATGTCCAAAAATTAACGCCG | 14701459 | 14701472 |
| SiSSM84994 | SisatBase | compound | 196 | AAACAGACGGGAGACGAAGA | GGGGAAAGGGCATAATGAAC | 14712805 | 14712854 |
| **SiSSM84996** | **SisatBase** | **compound** | **258** | **GGGAGGGAGACAGGGAAATA** | **ATTCGTGGTGATCGAGCTTT** | **14728083** | **14728148** |
| **SiSSM85023** | **SisatBase** | **(TG)10** | **262** | **GGCATGAGCCCACACATAAT** | **AAGACCGACAGCCGATTTTA** | **14765956** | **14765975** |

cont…

| **Marker ID** | **Source** | **SSR motif** | **Size** | **Primer Forward (5'-3')** | **Primer Reverse (5'-3')** | **Start (bp)** | **End (bp)** |
| --- | --- | --- | --- | --- | --- | --- | --- |
| SiSSM85041 | SisatBase | (GA)11 | 251 | ACGCCAAAATATTTGCCATA | TTCACCACAGCTCATGTTCC | 14812838 | 14812859 |
| SiSSM85056 | SisatBase | (AC)6 | 224 | TTGTTAGGAAATGGATCGGG | TGAGACGAGGGTGAGTGTTG | 14856950 | 14856961 |
| SiSSM85088 | SisatBase | (GT)9 | 262 | TCGGTGAATTTTGACTGACG | TCCTAAGCAAACGCAGTCCT | 14939936 | 14939953 |
| SiSSM85090 | SisatBase | (CT)9 | 135 | CATGACTTATGTATTGCCTATATGTGG | CGGATTGCAGTTTGTCGTTA | 14940947 | 14940964 |
| SiSSM85094 | SisatBase | (T)15(G)12 | 209 | TCCTTATGCCTTTGCACTCA | TCGCATAGGATCAAAATCACC | 14950201 | 14950227 |
| SiSSM85229 | SisatBase | (TA)17 | 240 | GAGGCACACAGAGGTCACAA | ACTCCTCTAATCCCCCTCCA | 15200463 | 15200496 |
| SiSSM85249 | SisatBase | (AT)19 | 278 | TCTCAATTATCAGAAGAAAGTCATAAA | CAAACCCACCTCATTAACCG | 15215174 | 15215211 |
| **SiSSM85354** | **SisatBase** | **(GA)15** | **161** | **TGTTCGCCTATTCTGCACTG** | **ACACACACATACACCCACGC** | **15489851** | **15489880** |
| **SiSSM85570** | **SisatBase** | **(GA)17** | **271** | **CTATCTGGGACGACGGAGAG** | **TTTCTTCAACAGATCCCGCT** | **15990137** | **15990170** |
| SiSSM85784 | SisatBase | (AT)12 | 255 | TGATGGCTTTGCTACTCACG | CTGAATTGCCCGTTGTACCT | 16364116 | 16364139 |
| 6390 | PMDBase | (AG)17 | 212 | GCTTCAGCCAGTTCTCCAAC | CAACCATATGTCGCTGTGCT | 16607555 | 16607588 |
| SiSSM86178 | SisatBase | (T)26 | 235 | AGCTATGCACATTTCCACCC | TGTACGTTCCTCGAGCACAA | 17182250 | 17182275 |
| SiSSM86181 | SisatBase | (TA)12 | 235 | TGACAAAAAGTGAACCAAAACAA | CATGACGTGGTTCAATGTCC | 17185785 | 17185808 |
| 6689 | PMDBase | (AT)9(AG)16 | 145 | TAATGGGCAGAGCTAATGGG | GCAAACATGAAGCCTTTGGT | 17212236 | 17212285 |
| SiSSM86323 | SisatBase | (AG)14 | 261 | AGCAAAGCACCACCTCAAGT | CCATCCTCTCGCTCTTCTTG | 17444408 | 17444435 |
| **SiSSM86906** | **SisatBase** | **(GA)23** | **221** | **AATCAAGATACGACGCAGGG** | **GTCCATTTCTCCTGGCCATA** | **18816333** | **18816378** |
| 7631 | PMDBase | (AT)22 | 193 | AGCACCTCTTGTGGTCCATC | TGCAATGAGTGGATTACCGA | 19392348 | 19392391 |

**^§^-** Polymorphic markers; **bold** letters indicate polymorphic markers between the two contrasting bulks and parents, Goenbaek (P1), Osan (P2), Milsung (P2)

**Supplemental Table S5.** Genomic information and annotation information of the SNPs from SSR flanking regions associated with *Phn-10* (*P.nicotianae* ) resistance.

| **Gene ID** | **Position** | **Physical position (Mb)^a^** | **Gene_annotation** |
| --- | --- | --- | --- |
| SIN_1019017 | 14538379 | 14,54 | ER Lumen Protein-Retaining Receptor |
| SIN_1019016 | 14554076 | 14,55 | NB-ARC-Probable Disease Resistance Protein At1g58390 |
| SIN_1019015 | 14560472 | 14,56 | NB-ARC-Putative Disease Resistance Protein At1g50180 |
| SIN_1019014 | 14570682 | 14,57 | Putative F-Box Protein At1g67623 |
| SIN_1019013 | 14574206 | 14,57 | Proline-Rich Receptor-Like Protein Kinase/Serine/Threonine-Protein Kinases |
| SIN_1019012 | 14583429 | 14,58 | Plasmodesmata Callose-Binding Protein 3 |
| SIN_1019011 | 14601277 | 14,60 | Protein Kinase Domain/Mitogen-Activated Protein Kinase Kinase Kinase YODA-Like |
| SIN_1019010 | 14603601 | 14,60 | AP2-ERF Domain/Ethylene-Responsive Transcription Factor ERF062 |
| SIN_1019009 | 14609126 | 14,61 | PREDICTED: Uncharacterized Protein |
| SIN_1019008 | 14630577 | 14,63 | ARID DNA-Binding Domain/High Mobility Group B Protein 15 |
| SIN_1019007 | 14634704 | 14,63 | DEAD-Box ATP-Dependent RNA Helicase 13 |
| SIN_1019006 | 14639219 | 14,64 | Repressed By RIM101 Protein 1 |
| SIN_1019005 | 14642509 | 14,64 | FYVE Zinc Finger-Myosin Heavy Chain, Non-Muscle-Like |
| SIN_1019004 | 14650492 | 14,65 | Pentatricopeptide (PRR) Repeat-Containing Protein At4g32430, Mitochondrial |
| SIN_1019003 | 14660753 | 14,66 | Ankyrin Repeat-Containing Domain |
| SIN_1019002 | 14666226 | 14,67 | Endoplasmic Reticulum (ER)-Associated Protein |
| SIN_1019001 | 14686769 | 14,69 | Serine-Threonine/Tyrosine-Protein Kinase, Catalytic Domain |
| SIN_1019000 | 14692204 | 14,69 | Homeobox-Like Domain Superfamily |
| SIN_1018999 | 14700937 | 14,70 | F-Box Protein PP2-B5/ Phloem Protein 2-Like |
| SIN_1018998 | 14706989 | 14,71 | Serine/Threonine-Protein Phosphatase PP1 Tetraphosphatase |
| SIN_1018997 | 14724989 | 14,72 | F-Box Protein PP2-B5/ |
| SIN_1018996 | 14727979 | 14,73 | F-Box Protein PP2-B5/ Phloem Protein 2-Like |
| SIN_1018995 | 14734025 | 14,73 | F-Box Protein PP2-B5/ Phloem Protein 2-Like |
| SIN_1018994 | 14737451 | 14,74 | Probable Pectinesterase 29, Catalytic /Cell Wall |
| SIN_1018993 | 14751096 | 14,75 | 60S Ribosomal Protein L13a-4-Like |
| SIN_1018992 | 14766489 | 14,77 | Myc-Type, Basic Helix-Loop-Helix (Bhlh) Domain |

**Supplemental Table S6.** The sequence matched with candidate genes using BLAST and *S. indicum* (v.2) reference genome for identification of putative genes within associated genomic region determined by GWAS and Haplotype analysis.

| **SNP** | **Haplotype** | | | | | | **Gene ID** | **Position** | | **Description** | |
| --- | --- | --- | --- | --- | --- | --- | --- | --- | --- | --- | --- |
| S10_14456135  *(No2040) S10_14532795 (No2040)*  S10_14563939 *(KACC48120, KACC48121, and No2040)*  S10_14643881 *(No2040)* S10_14665280 *(No2040)* S10_14693190 *(No2040)* | **28 genes (244.0 Kb)** |  | |  | |  | SIN_1019027 | 14451847..14466672 (- strand) | | Serine/Threonine Protein Phosphatase 2A Regulatory Subunit B''beta | |
|  |  |  | |  | |  | SIN_1019026 | 14469278..14478285 (- strand) | | Ubiquitin Carboxyl-Terminal Hydrolase 2 | |
|  |  |  | |  | |  | SIN_1019025 | 14483080..14483415 (+ strand) | | Uncharacterized Protein LOC110011297 | |
|  |  |  | |  | |  | SIN_1019024 | 14484966..14487029 (+ strand) | | Elongation Factor 1-Alpha-Like / Hypothetical Protein CDL15_Pgr022686 | |
|  |  |  | |  | |  | SIN_1019023 | 14489808..14493423 (+ strand) | | Putative Disease Resistance Protein At1g59780 | |
|  |  |  | |  | |  | SIN_1019022 | 14509311..14512398 (+ strand) | | Disease Resistance Protein RPP8-Like | |
|  |  |  | |  | |  | SIN_1019021 | 14520679..14525469 (+ strand) | | Small Ubiquitin-Related Modifier 2-Like | |
|  |  |  | |  | |  | SIN_1019020 | 14528970..14531008 (+ strand) | | Small Ubiquitin-Related Modifier 1 | |
|  |  |  | |  | |  | SIN_1019019 | 14532148..14533208 (+ strand) | | Cytochrome P450 71D95-Like / Hypothetical Protein MIMGU_Mgv1a025918mg | |
|  |  |  | |  | |  | SIN_1019018 | 14534978..14535630 (+ strand) | | Small Ubiquitin-Related Modifier 2 | |
|  |  |  | |  | |  | SIN_1019017 | 14538379..14542560 (- strand) | | ER Lumen Protein-Retaining Receptor | |
|  |  |  | |  | |  | SIN_1019016 | 14554076..14557285 (- strand) | | **Probable Disease Resistance Protein At1g58390** | |
|  |  | **38 genes ( 301 Kb)** | | **56 genes ( 434 Kb)** | |  | SIN_1019015 | 14560472..14564212 (- strand) | | Putative Disease Resistance Protein At1g50180 | |
|  |  |  |  |  |  |  | SIN_1019014 | 14570682..14570930 (+ strand) | | Putative F-Box Protein At1g67623 | |
|  |  |  |  |  |  |  | SIN_1019013 | 14574206..14579023 (- strand) | | **Proline-Rich Receptor-Like Protein Kinase PERK3** | |
|  |  |  |  |  |  |  | SIN_1019012 | 14583429..14587250 (- strand) | | Plasmodesmata Callose-Binding Protein 3 | |
|  |  |  |  |  |  |  | SIN_1019011 | 14601277..14601633 (+ strand) | | Mitogen-Activated Protein Kinase Kinase Kinase YODA-Like | |
|  |  |  |  |  |  |  | SIN_1019010 | 14603601..14604818 (+ strand) | | Ethylene-Responsive Transcription Factor ERF062 | |
|  |  |  |  |  |  |  | SIN_1019009 | 14609126..14609542 (- strand) | | Uncharacterized Protein | |
|  |  |  |  |  |  |  | SIN_1019008 | 14630577..14634595 (- strand) | | High Mobility Group B Protein 15 | |
|  |  |  |  |  |  |  | SIN_1019007 | 14634704..14635027 (- strand) | | Unknown | |
|  |  |  |  |  |  |  | SIN_1019006 | 14639219..14641353 (- strand) | | Repressed By RIM101 Protein 1 | |
|  |  |  |  |  |  |  | SIN_1019005 | 14642509..14648243 (+ strand) | | Myosin Heavy Chain, Non-Muscle-Like | |
|  |  |  |  |  |  |  | SIN_1019004 | 14650492..14654632 (- strand) | | Pentatricopeptide Repeat-Containing Protein At4g32430 | |
|  |  |  |  |  |  |  | SIN_1019003 | 14660753..14665846 (+ strand) | | Uncharacterized Protein | |
|  |  |  |  |  |  |  | SIN_1019002 | 14666226..14672207 (- strand) | | Uncharacterized Protein | |
|  |  |  |  |  |  |  | SIN_1019001 | 14686769..14689240 (+ strand) | | **Receptor Like Protein Kinase S.2** | |
|  |  |  |  |  |  |  | SIN_1019000 | 14692204..14695873 (- strand) | | Protein PHR1-LIKE 2 | |
| S10_14563939 *(KACC48120, KACC48121, No2040)*  S10_14830583 *(KACC48120)* S10_14830584 *(KACC48120)* S10_14830631*(KACC48120)*  S10_14830687 *(KACC48120, KACC48121*)  S10_14841897 *(KACC48120)* S10_14860890 *(KACC48120, KACC48121)*  S10_14860891 *(KACC48120, KACC48121)* S10_14860929 *(KACC48120, KACC48121)*  S10_14860930 *(KACC48120, KACC48121)*  S10_14864663 *(KACC48121)* S10_14877987 *(KACC48121)* S10_14965952 *(KACC48121)* S10_14990956 *(KACC48121)* |  |  |  |  |  |  | SIN_1018999 | 14700937..14705072 (+ strand) | | F-Box Protein PP2-B5 | |
|  |  |  |  |  |  |  | SIN_1018998 | 14706989..14711116 (+ strand) | | Serine/Threonine-Protein Phosphatase PP1 | |
|  |  |  |  |  |  |  | SIN_1018997 | 14724989..14727081 (+ strand) | | F-Box Protein PP2-B5 | |
|  |  |  |  |  |  |  | SIN_1018996 | 14727979..14730530 (+ strand) | | F-Box Protein PP2-B5 | |
|  |  |  |  |  |  |  | SIN_1018995 | 14734025..14736558 (+ strand) | | F-Box Protein PP2-B5 | |
|  |  |  |  |  |  |  | SIN_1018994 | 14737451..14746414 (- strand) | | Probable Pectinesterase 29 | |
|  |  |  |  |  |  |  | SIN_1018993 | 14751096..14752984 (- strand) | | 60S Ribosomal Protein L13a-4-Like | |
|  |  |  |  |  |  |  | SIN_1018992 | 14766489..14767850 (- strand) | | Transcription Factor FAMA | |
|  |  |  |  |  |  |  | SIN_1018991 | 14771856..14772131 (+ strand) | | Unknown | |
|  |  |  |  |  |  |  | SIN_1018990 | 14774536..14776227 (- strand) | | **Putative Disease Resistance RPP13-Like Protein 1** | |
|  |  |  |  |  |  |  | SIN_1018989 | 14778561..14779476 (+ strand) | | Uncharacterized Protein | |
|  |  |  |  |  |  |  | SIN_1018988 | 14785577..14787307 (- strand) | | Putative Disease Resistance Protein RGA3 | |
|  |  |  |  |  |  |  | SIN_1018987 | 14789609..14790757 (+ strand) | | Uncharacterized LOC110012745 | |
|  |  |  |  |  |  |  | SIN_1018986 | 14793789..14796495 (+ strand) | | Cytochrome B5 | |
|  |  |  |  |  |  |  | SIN_1018985 | 14797190..14799002 (- strand) | | Cyclin-Dependent Kinase Inhibitor 3 | |
|  |  |  |  |  |  |  | SIN_1018984 | 14803602..14805582 (+ strand) | | Pentatricopeptide Repeat-Containing Protein At4g13650-Like | |
|  |  |  |  |  |  |  | SIN_1018983 | 14811413..14811823 (+ strand) | | Unknown | |
|  |  |  |  |  |  |  | SIN_1018982 | 14817324..14819102 (+ strand) | | Unknown | |
|  |  |  |  |  |  |  | SIN_1018981 | 14820862..14821703 (+ strand) | | Unknown | |
|  |  |  |  |  |  |  | SIN_1018980 | 14830448..14838296 (+ strand) | | Unnamed Protein Product | |
|  |  |  |  |  |  |  | SIN_1018979 | 14839170..14845554 (- strand) | | Probable Disease Resistance Protein RF45 | |
|  |  |  |  |  |  |  | SIN_1018978 | 14847198..14865712 (- strand) | | Probable Disease Resistance Protein RF45 | |
| S10_14880164 *(KACC48120, KACC48121)*  S10_14925756 *(KACC48120)* S10_14925898 *(KACC48120)* S10_14925982 *(KACC48120, KACC48121)*  S10_14925999 *(KACC48120, KACC48121)* S10_14927344 *(KACC48120, KACC48121)*  S10_14927412 *(KACC48120, KACC48121)* S10_14927451 *(KACC48120, KACC48121)*  S10_14934271 *(KACC48120, KACC48121)*  S10_14934878 *(KACC48120, KACC48121)*  S10_14972452 *(KACC48120, KACC48121)* S10_15011585 *(KACC48120, KACC48121)*  S10_15019593 *(KACC48120)* |  | **18 genes ( 133 Kb)** | |  |  |  | SIN_1018975 | 14877429..14880295 (+ strand) | | Cytochrome P450 71A1-Like | |
|  |  |  |  |  |  |  | SIN_1018974 | 14882704..14884595 (+ strand) | | Cytochrome P450 71A1-Like | |
|  |  |  |  |  |  |  | SIN_1018973 | 14888800..14893174 (+ strand) | | Uncharacterized Protein | |
|  |  |  |  |  |  |  | SIN_1018972 | 14903157..14904581 (+ strand) | | Hydroquinone Glucosyltransferase-Like | |
|  |  |  |  |  |  |  | SIN_1018971 | 14906134..14910502 (+ strand) | | Cellulose Synthase-Like Protein D1 | |
|  |  |  |  |  |  |  | SIN_1018970 | 14916358..14918488 (- strand) | | **Vesicle-Associated Membrane Protein 722** | |
|  |  |  |  |  |  |  | SIN_1018969 | 14920924..14923849 (- strand) | | Ribonuclease P Protein Subunit P25-Like Protein | |
|  |  |  |  |  |  |  | SIN_1018968 | 14924429..14924668 (- strand) | | Caffeoylshikimate Esterase | |
|  |  |  |  |  |  |  | SIN_1018967 | 14926043..14928179 (+ strand) | | Uncharacterized Aarf Domain-Containing Protein Kinase At1g79600, Chloroplastic | |
|  |  |  |  |  |  |  | SIN_1018966 | 14929389..14935720 (+ strand) | | Uncharacterized Aarf Domain-Containing Protein Kinase At1g79600, Chloroplastic | |
|  |  |  |  |  |  |  | SIN_1018965 | 14936297..14937565 (- strand) | | Pentatricopeptide Repeat-Containing Protein At3g26630, Chloroplastic | |
|  |  |  |  |  |  |  | SIN_1018964 | 14940852..14946343 (+ strand) | | Ankyrin Repeat Domain-Containing Protein 13C-A-Like | |
|  |  |  |  |  |  |  | SIN_1018963 | 14947161..14947986 (- strand) | | Uncharacterized Protein TCM_020382 | |
|  |  |  |  |  |  |  | SIN_1018962 | 14958873..14966719 (+ strand) | | Ubiquinone Biosynthesis Monooxygenase COQ6, Mitochondrial | |
|  |  |  |  |  |  |  | SIN_1018961 | 14971538..14973984 (- strand) | | Chalcone Synthase | |
|  |  |  |  |  |  |  | SIN_1018960 | 14990068..14991794 (- strand) | | Chalcone Synthase J-Like | |
|  |  |  |  |  |  |  | SIN_1018959 | 15004626..15007378 (- strand) | | Chalcone Synthase-Like | |
|  |  |  |  |  |  |  | SIN_1018958 | 15011374..15013327 (+ strand) | | E3 Ubiquitin-Protein Ligase RLIM | |
| S10_15031344 *(KACC48120, KACC48121, and No2040)* S10_15081876 *(KACC48120, KACC48121, and No2040)* S10_15091261 *(KACC48120, KACC48121, and No2040)* S10_15098669 *(KACC48120, KACC48121, and No2040)* S10_15109810 *(KACC48120, KACC48121, and No2040)* S10_15109872 *(KACC48120, KACC48121, and No2040)* S10_15109921 *(KACC48120, KACC48121, and No2040)* | | | **12 genes (79 Kb)** | | SIN_1018957 | | | | 15027416..15032237 (- strand) | | 3-Ketoacyl-Coa Thiolase 2, Peroxisomal |
|  |  |  |  |  | SIN_1018956 | | | | 15040719..15046302 (- strand) | | Mitosis Inhibitor Protein Kinase Wee1 |
|  |  |  |  |  | SIN_1018955 | | | | 15048690..15052668 (- strand) | | Probable Voltage-Gated Potassium Channel Subunit Beta |
|  |  |  |  |  | SIN_1018954 | | | | 15057543..15057926 (+ strand) | | Uncharacterized Protein |
|  |  |  |  |  | SIN_1018953 | | | | 15060887..15061297 (+ strand) | | Uncharacterized Protein |
|  |  |  |  |  | SIN_1018952 | | | | 15082220..15086159 (+ strand) | | Probable Pectate Lyase 8 |
|  |  |  |  |  | SIN_1018951 | | | | 15091397..15092469 (+ strand) | | Acanthoscurrin-1 |
|  |  |  |  |  | SIN_1018950 | | | | 15094456..15094857 (+ strand) | | Unknown |
|  |  |  |  |  | SIN_1018949 | | | | 15098454..15098759 (- strand) | | Auxin-Induced Protein 15A-Like |
|  |  |  |  |  | SIN_1018948 | | | | 15102253..15102654 (+ strand) | | Uncharacterized Protein |
|  |  |  |  |  | SIN_1018947 | | | | 15103594..15104126 (+ strand) | | Hypothetical Protein F511_02833 |
|  |  |  |  |  | SIN_1018946 | | | | 15106078..15106702 (+ strand) | | Hypothetical Protein F511_02833 |

**Bold and underlines** are designed gene-specific primers

**Supplemental Table S7**. List of oligonucleotides used for qRT-PCR analyses.

| **oligo name** | **5′-3′primer sequence** | **3′-5′primer sequence** | **Product size** | **Gene name** |
| --- | --- | --- | --- | --- |
| Phn-10-1-qPCR | GGGCACAGAGTTGTTTCCAT | GATGGCGGTCCAGCTTAATA | 210 | LOC105171879 receptor like protein kinase S.2 |
| Phn-10-2-qPCR | TTTCCCGCTTAACAGGATTG | AAGTTGCCAAACCATTCAGG | 232 | LOC105171888 proline-rich receptor-like protein kinase PERK3 |
| Phn-10-3-qPCR | CCTGAATGGTTTGGCAACTT | ACGGGATATGCGAAATCTTG | 214 | LOC105172053 putative disease resistance RPP13-like protein 1 |
| Phn-10-4-qPCR | GCAGAAAATTGGCGATGATT | ACAATGAATGGAGGGCAAAG | 239 | **LOC110012696 probable disease resistance protein At1g58390** |
| Phn-10-5-qPCR | TGAAACTGCCGTAACTAGCATC | TTCCTCACCTGTGTCCCTTT | 175 | LOC105171860 vesicle-associated membrane protein 722 |
| Si18SrRNA-qPCR | CGTCCCTGCCCTTTGTACAC' | CGAACACTTCACCGGACCAT' |  | as actin reference gene |
